# Glycan-costumed virus-like particles promote type 1 anti-tumor immunity

**DOI:** 10.1101/2024.01.18.575711

**Authors:** Valerie Lensch, Adele Gabba, Robert Hincapie, Sachin H. Bhagchandani, Ankit Basak, Mohammad Murshid Alam, Darrell J. Irvine, Alex K. Shalek, Jeremiah A. Johnson, M. G. Finn, Laura L. Kiessling

**Author notes:** These authors contributed equally to this work (Adele Gabba, Robert Hincapie).

## Abstract

Cancer vaccine development is inhibited by a lack of strategies for directing dendritic cell (DC) induction of effective tumor-specific cellular immunity. Pathogen engagement of DC lectins and toll-like receptors (TLRs) shapes immunity by directing T cell function. Strategies to activate specific DC signaling pathways via targeted receptor engagement are crucial to unlocking type 1 cellular immunity. Here, we engineered a glycan-costumed virus-like particle (VLP) vaccine that delivers programmable peptide antigens to induce tumor-specific cellular immunity *in vivo*. VLPs encapsulating TLR7 agonists and decorated with a selective mannose-derived ligand for the lectin DC-SIGN induced robust DC activation and type 1 cellular immunity, whereas VLPs lacking this key DC-SIGN ligand failed to promote DC-mediated immunity. Vaccination with glycan-costumed VLPs generated tumor antigen-specific Th1 CD4^+^ and CD8^+^ T cells that infiltrated solid tumors, inhibiting tumor growth in a murine melanoma model. Thus, VLPs employing lectin-driven immune reprogramming provide a framework for advancing cancer immunotherapies.

## Introduction

Vaccines have transformed the fight against critical infectious diseases; however, developing vaccines against cancer has proved challenging.^1^ Therapeutic cancer vaccines require type 1 cellular immunity—both helper type 1 (Th1 CD4^+^) and cytotoxic (CD8^+^) T cells that recognize and kill cancer cells;^2^ therefore, strategies to induce tumor-specific type 1 T cells *in vivo* are needed. Dendritic cells (DCs) are the immune system’s central player for initiating anti-tumor immunity.^3^ As professional antigen-presenting cells (APCs), DCs internalize pathogens and present processed epitopes on major histocompatibility complex (MHC) molecules for T cell priming.^4^ Additionally, in response to pathogen engagement, DCs secrete cytokines that activate cellular (Th1 and CD8^+^), humoral (Th2), or tolerogenic (Treg) T cell immunity.^5^ Due to their key role in modulating immunity via T cell polarization, specific activation of individual DC signaling pathways is critical for inducing tumor antigen-specific, type 1 cellular immunity.

DC-based immunotherapies primarily rely on *ex vivo* activation and engineering of DCs to present tumor antigens for T cell priming.^6^ *Ex vivo* loaded DCs are compromised by limited migration to lymph nodes for T cell interactions and a diminished capacity to maintain pro-inflammatory status upon injection, which can inadvertently lead to the undesired induction of Treg and Th2 responses.^6^ In contrast, targeting of antigens to DCs *in vivo* could enable the direct activation of DCs for generation of pro-inflammatory, tumor-specific immune responses. This approach would also circumvent the resource-intensive *ex vivo* DC generation process to enable scalable vaccine production.

DCs employ lectin recognition of cell surface glycans as the first extracellular determination of self and non-self. DC lectins can identify pathogen-associated glycans as foreign threats to ramp up immunity and self-associated glycans as signals to induce tolerance, preventing autoimmunity.^7^ However, aberrant glycosylation exhibited by cancer cells often suppresses immunity through inhibitory lectin signaling, notably exemplified by the sialic acid-Siglec axis.^8^ The ability of lectins to shape DC signaling and modulate downstream immunity make them interesting targets for immunomodulation.^9, 10^ The dendritic cell-specific surface lectin DC-SIGN (CD209) binds mannose or fucose-substituted oligosaccharides on numerous exogenous pathogens with distinct immune outcomes.^11-13^ While DC-SIGN binding facilitates efficient pathogen uptake, engagement of DC-SIGN alone may induce regulatory responses and immune evasion.^14-16^ Lectins are therefore pivotal regulators of DC immunity, but they do not act alone.

Toll-like receptors (TLRs) bind a broader range of viral and bacterial molecular structures determined by their location on the cell surface or within endosomes of DCs. The endosomal TLR7 binds viral and microbial nucleic acids, increasing formation of MHC-peptide complexes, upregulation of costimulatory molecules (CD80, CD86, CD40), and secretion of IL-12.^17-19^ TLR7 agonists and TLR combinations have been studied as vaccine adjuvants and cancer therapeutics,^20-23^ but challenges in achieving suitable efficacy while avoiding toxicity remain a significant barrier to their use.^24^ While nonselective lectin-targeting has been used to facilitate uptake of TLR agonists by APCs, the reported results have varying degrees of success, potentially due to poorly understood immune outcomes of engaging both lectins and TLRs.^25-29^ As co-delivery of ligands for lectins and TLRs has the potential to switch immune outcomes from regulatory to immunostimulatory,^30^ we envisaged that selective dual lectin-TLR signaling could be leveraged to activate immunity. We recently reported an aryl mannoside ligand (Man) that selectively binds to DC-SIGN and found that synergistic engagement of DC-SIGN and TLR7 by Man-functionalized virus-like particles (VLPs) resulted in Th1 signaling.^31^ The chemical structure of the mannose ligand was critical for DC-SIGN signaling. Man bound DC-SIGN with higher avidity compared to a trimannose ligand at lower pH, a feature designed to afford enhanced engagement of DC-SIGN and TLR7 in the endosome for robust DC activation. Given that type 1 cellular immunity is critical for therapeutic cancer vaccine efficacy, we envisioned that delivering tumor antigens within this framework could generate tumor-specific CD8^+^ T cells and robustly reprogram the immune system’s response to cancer.

Here, we leverage lectin-driven immune reprogramming to induce potent anti-tumor immunity. Our approach involves engineering glycan-costumed VLPs that multivalently display synthetic DC-SIGN ligands and tumor antigens as well as package single-stranded RNA (ssRNA) TLR7 agonists to induce type 1 cellular immunity *in vivo* (Fig. 1a,b). The mannosylated VLPs exhibited efficient and selective internalization via DC-SIGN, while simultaneous engagement of DC-SIGN and TLR7 resulted in robust DC activation and the generation of effective tumor antigen-specific T cells for improved tumor clearance. The platform reported here was shown to be an effective, self-adjuvanting vaccine system that leverages DC-SIGN binding to generate robust antigen-specific type 1 cellular immunity.

**Fig. 1.**
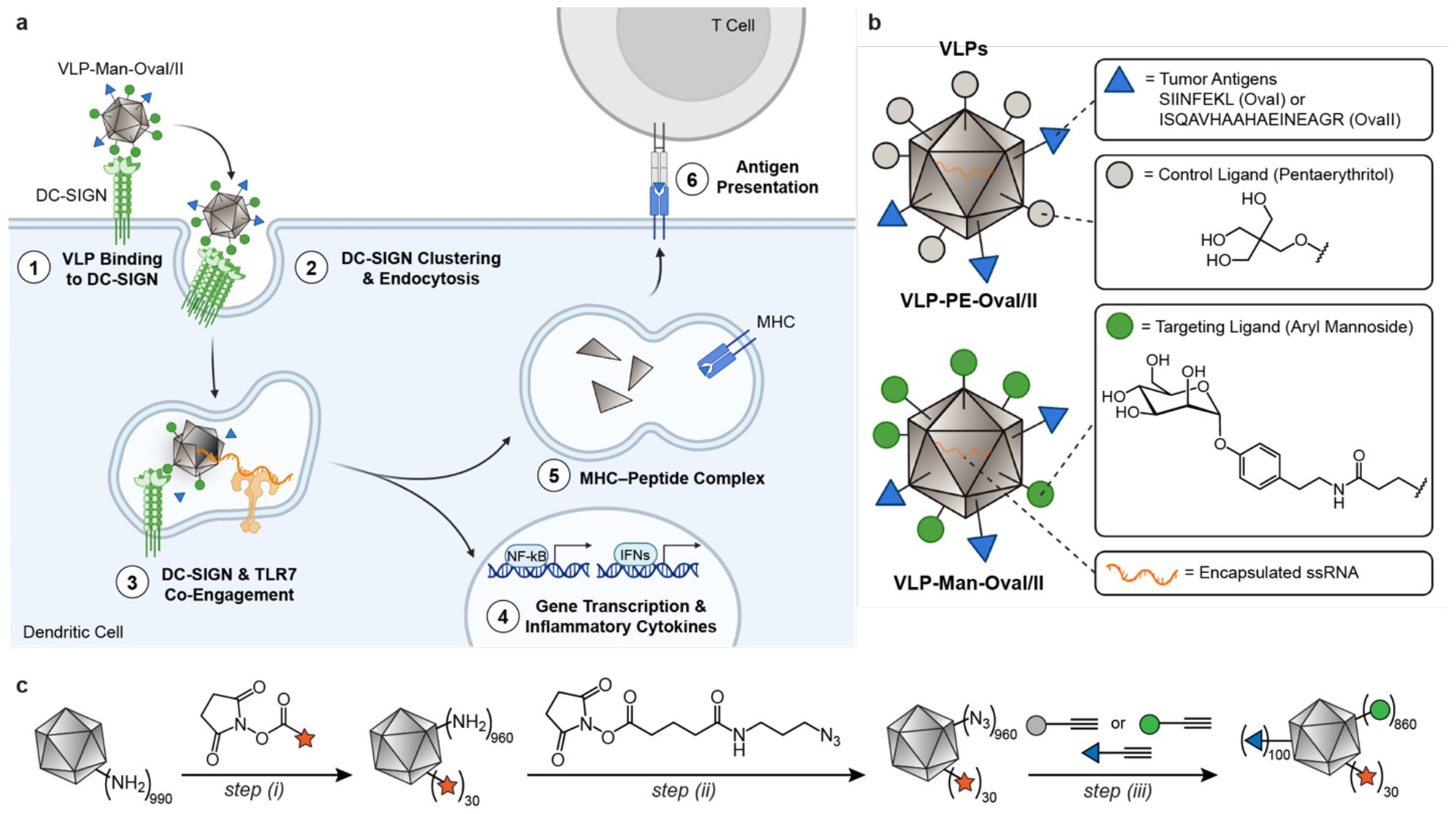
Aryl mannoside-substituted VLPs as a platform for DC-targeted cancer vaccines. **a**, Schematic of VLP-Man-OvaI/II engagement of DC-SIGN and TLR7 on DCs for antigen presentation and T cell priming. VLP decoration with the aryl mannoside ligand facilitates DC targeting and antigen internalization. Following endocytosis, aryl mannoside-substituted VLPs maintain DC-SIGN binding and facilitate ssRNA engagement of endosomal TLR7 to activate DCs and induce tumor antigenspecific T cells. **b**, Design of VLPs surface-functionalized with tumor antigens OvaI and OvaII, control pentaerythritol (PE) or Man ligands, and encapsulated ssRNAs. **c**, Surface accessible lysine residues were functionalized to incorporate an azide handle and then subjected to a copper(I)-catalyzed Huisgen azide–alkyne cycloaddition (CuAAC), a type of click chemistry, to append tumor antigens and ligands. Steps (i, ii): 0.1 M potassium phosphate buffer, pH 7.4; room temperature, 2 h; step (iii): tris(3 hydroxypropyltriazolylmethyl)amine (THPTA), CuSO_4_, sodium ascorbate, aminoguanidine, 0.1 M potassium phosphate, pH 7.0, 50 °C, 1 h.

### Preparation and functionalization of VLPs

Given that specific lectin-TLR dual activation is poorly understood, we sought to engineer VLPs that could specifically engage DC-SIGN and TLR7 to elicit robust DC activation. VLPs are non-replicating nanostructures comprised of self-assembling virus-derived coat protein monomers.^32^ Encapsulation of single-stranded RNAs (ssRNAs) from the bacterial expression host enable TLR7 signaling for immune activation.^33^ VLPs have been successfully used to induce humoral immunity comprising long-lasting IgG antibody responses,^34-39^ but engineering VLPs to induce effective cellular immunity remains challenging. We previously reported that VLPs engineered from bacteriophage Qβ to present a DC-SIGN-selective, pH-independent aryl mannoside ligand and bearing genetically encoded OVA(323-339) (OvaII) peptides could induce antigen-specific Th1 CD4^+^ T cells *in vivo*.^31^ Motivated by these results, we hypothesized that tumor antigen delivery in this context could induce both tumor antigen-specific Th1 CD4^+^ and CD8^+^ T cells, providing holistic type 1 cellular immunity. However, analogous Qβ VLPs bearing a similar density of Man and genetically encoded OVA(257-264) (OvaI) peptides towards induction of antigen-specific CD8^+^ T cells were not accessible, as the internal lysine residue of OvaI would likely be acylated in the course of particle conjugation. Attempts to chemically derivatize Qβ VLPs with high densities of glycans and peptides (>540 Man and >40 peptides per particle) were not successful, since these particles were prone to irreversible aggregation.

To maximize the loading of peptide antigens on VLPs, we turned to the bacteriophage PP7-PP7 platform, which has a higher density of accessible amines for functionalization (∼1200 per particle), and a higher tolerance for peptide incorporation.^40^ To enable particle loading with diverse peptide antigens, including peptides containing nucleophilic amino acids (*e*.*g*., lysine, cysteine), we designed a two-stage modification strategy. Amine-reactive esters were used to install a low density of fluorophores and a high density of azides on VLPs. The alkyne-containing aryl mannoside ligand was synthesized by a facile, three-step synthesis (Supplementary Scheme 1). Following particle purification, the intermediate particles were then coupled to alkyne-containing glycomimetic or control ligands (Man, PE) and peptide antigens (OvaI, OvaII) via a copper(I)-catalyzed Huisgen azide–alkyne cycloaddition (CuAAC), a powerful click reaction (Fig. 1c).^41, 42^ The model tumor antigens OvaI and OvaII were modified with a cathepsin D-sensitive linker to facilitate release by endosomal proteolytic processing and loading on MHC molecules.^43^ The extent of ligand and antigen functionalization on each particle was controlled by sequential addition of peptide-alkynes and Man-/PE-alkynes. The resulting conjugates were analyzed by LC-ESI-TOF-MS to determine the average loading density of each functional unit (Supplementary Fig. 1) and by DLS and FPLC to confirm particle integrity (Supplementary Fig. 2,3), demonstrating that all particles displayed equivalent ligand loadings and size (∼50 nm diameter). Thus, we generated PP7 particles displaying PE_1000_ (VLP-PE), PE_900_-OvaI_100_ (VLP-PE-OvaI), PE_900_-OvaII_100_ (VLP-PE-OvaII), Man_1000_ (VLP-Man), Man_900_-OvaI_100_ (VLP-Man-OvaI), and Man_900_-OvaII_100_ (VLP-Man-OvaII) (Supplementary Table 1,2). We hypothesized that particles constructed in this manner could enhance DC activation via costimulatory signaling by dual mannose-DC-SIGN and ssRNA-TLR7 interactions to achieve antigen-selective immunity. Additionally, we postulated that combinations of particles delivering MHC-I and MHC-II epitopes (*e*.*g*., VLP-Man-OvaI/II) could induce both CD4^+^ and CD8^+^ T cells, where Th1 CD4^+^ T cells could help augment CD8^+^ T cell proliferation, for complete and robust anti-tumor immunity.

### Engineered VLPs engage DC-SIGN and TLR7 for efficient uptake and DC activation

Receptor recognition and signaling facilitates DC activation and antigen uptake for efficient T cell priming.^4^ To investigate the efficiency of DC-SIGN-mediated particle uptake and DC activation, we compared the uptake of mannosylated VLPs against control particles. Fluorophore-labeled VLPs were added to monocyte-derived dendritic cells (moDCs), and VLP uptake was monitored by flow cytometry. DC uptake of VLP-Man-OvaI/II was significantly higher (5-fold) than that of VLP-PE-OvaI/II over 30 min (Fig. 2a,b), demonstrating that conjugation of tumor antigens did not interfere with the glycan-lectin interactions. To investigate the endocytic role of DC-SIGN over other mannose-binding lectins, moDCs were pre-treated with blocking antibodies against DC-SIGN, mannose receptor (MR), Dectin-2, or Mincle and then exposed to VLPs. Pre-treatment with DC-SIGN—but not MR, Dectin-2, or Mincle— resulted in a significant decrease in VLP-Man-OvaI/II uptake, confirming that DC-SIGN is the major endocytic receptor for VLP-Man-OvaI/II (Fig. 2c). These data indicate that VLP-Man-OvaI/II uptake occurs primarily by DC-SIGN-mediated endocytosis.

**Fig. 2.**
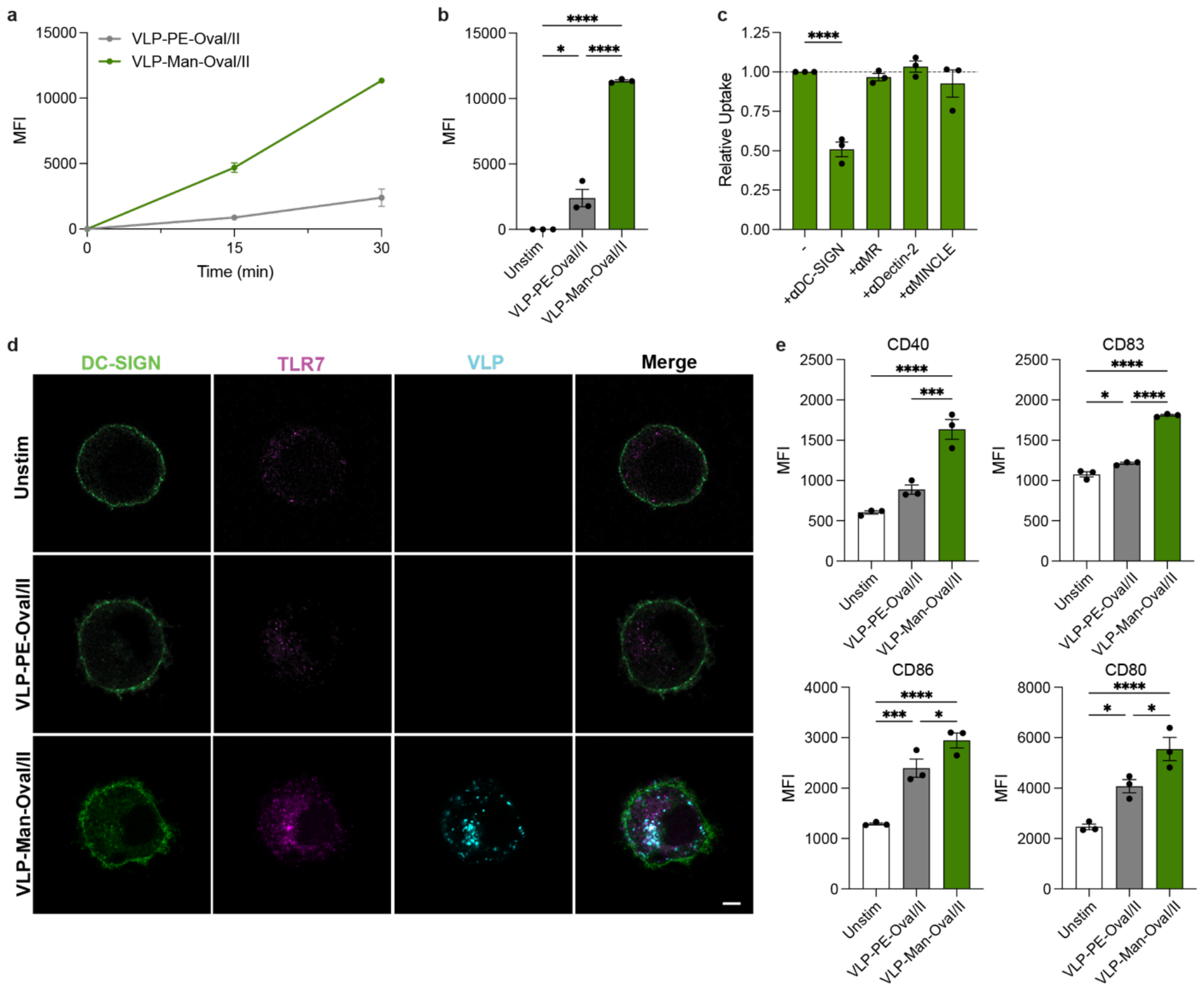
VLP-Man-OvaI/II co-engages DC-SIGN and TLR7 for DC activation. **a**, moDCs were incubated with 8 nM AF647-labeled VLPs for 30 min, and uptake of VLP by moDCs was measured by flow cytometry. **b**, Mean fluorescence intensity of AF647-labeled VLPs in CD11c^+^ moDCs indicated at 30 min. **c**, moDCs were pre-treated with blocking antibodies against lectins for 20 min and then incubated with 8 nM VLP-Man-OvaI/II for 15 min. The uptake of VLPs was assessed by flow cytometry. Relative uptake was calculated by normalization against a control sample without pre-treatment of blocking antibodies. **d**, Confocal microscopy images showing VLP localization in moDCs after 1 h incubation with 8 nM VLPs. Green: AF405 anti-DC-SIGN; Pink: AF488 anti-TLR7; Cyan: AF647-VLP. Scale bars, 5 μm. **e**, Activation of moDCs by VLPs. moDCs were incubated with 8 nM VLPs for 24 h, and the expression levels of activation surface markers CD40, CD83, CD86, and CD80 were measured by flow cytometry. Data represent mean +/-standard error of the mean (s.e.m.) from a representative experiment (n=3). Statistical analysis was performed using one-way ANOVA with Bonferroni’s multiple comparisons test. *P<0.1, **P<0.01, ***P<0.001, ****P<0.0001. Each experiment was repeated at least twice with similar results.

We envisioned that the endocytic ability of DC-SIGN would facilitate delivery of VLP-encapsulated ssRNAs to endosomal TLR7.^44^ To assess co-engagement of DC-SIGN and TLR7, we performed confocal microscopy on moDCs stimulated with VLPs and labeled with anti-DC-SIGN and anti-TLR7 antibodies. Confocal images revealed that VLP-Man-OvaI/II was efficiently endocytosed by moDCs within 30 min whereas VLP-PE-OvaI/II was present only at low levels in the cells, corroborating the enhanced uptake of mannosylated VLPs as measured by flow cytometry. Additionally, colocalization of only VLP-Man-OvaI/II with both DC-SIGN and TLR7 was observed (Fig. 2d), further confirming DC-SIGN and TLR7 engagement by VLP-Man-OvaI/II.

To assess DC activation by VLPs, moDCs were incubated with VLP-Man-OvaI/II or VLP-PE-OvaI/II for 24 h. VLP-Man-OvaI/II induced more robust DC activation than VLP-PE-OvaI/II, as demonstrated by the significant increase in surface expression of activation markers and costimulatory molecules CD40, CD80, CD83, and CD86 (Fig. 2e). Taken together, the mannosylated VLPs target DCs via binding to the cell surface lectin DC-SIGN, facilitating highly efficient endocytosis and engagement of TLR7 for robust DC activation.

### VLP-Man elicits type 1 immune responses in vitro

To better understand the basis for the enhanced upregulation of activation markers observed upon stimulation with mannosylated VLPs, we performed bulk RNA-seq experiments on moDCs treated with VLP-Man-OvaI/II and VLP-PE-OvaI/II or unstimulated control moDCs. A total of 486 genes were differentially expressed after treatment with VLP-Man-OvaI/II vs. VLP-PE-OvaI/II (log_2_FC >1.5 and p<0.05) (Fig. 3a). The significantly upregulated genes included *CXCL1, CXCL5, ISG15, IL6, CCL4, ISG20*, and *SOCS3*, all of which are implicated in the TLR7 downstream signaling pathway,^45^ suggesting a higher extent of TLR7 activation with the mannosylated VLPs (Fig. 3b). Gene set enrichment analysis (GSEA) between VLP-Man-OvaI/II- and VLP-PE-OvaI/II-treated moDCs revealed that hallmark DC activation pathways were also upregulated in VLP-Man-OvaI/II-treated DCs, including pathways related to IFN-α, IFN-γ, and TNF-α signaling via nuclear factor-κB (NF-kB) (Fig. 3c).

**Fig. 3.**
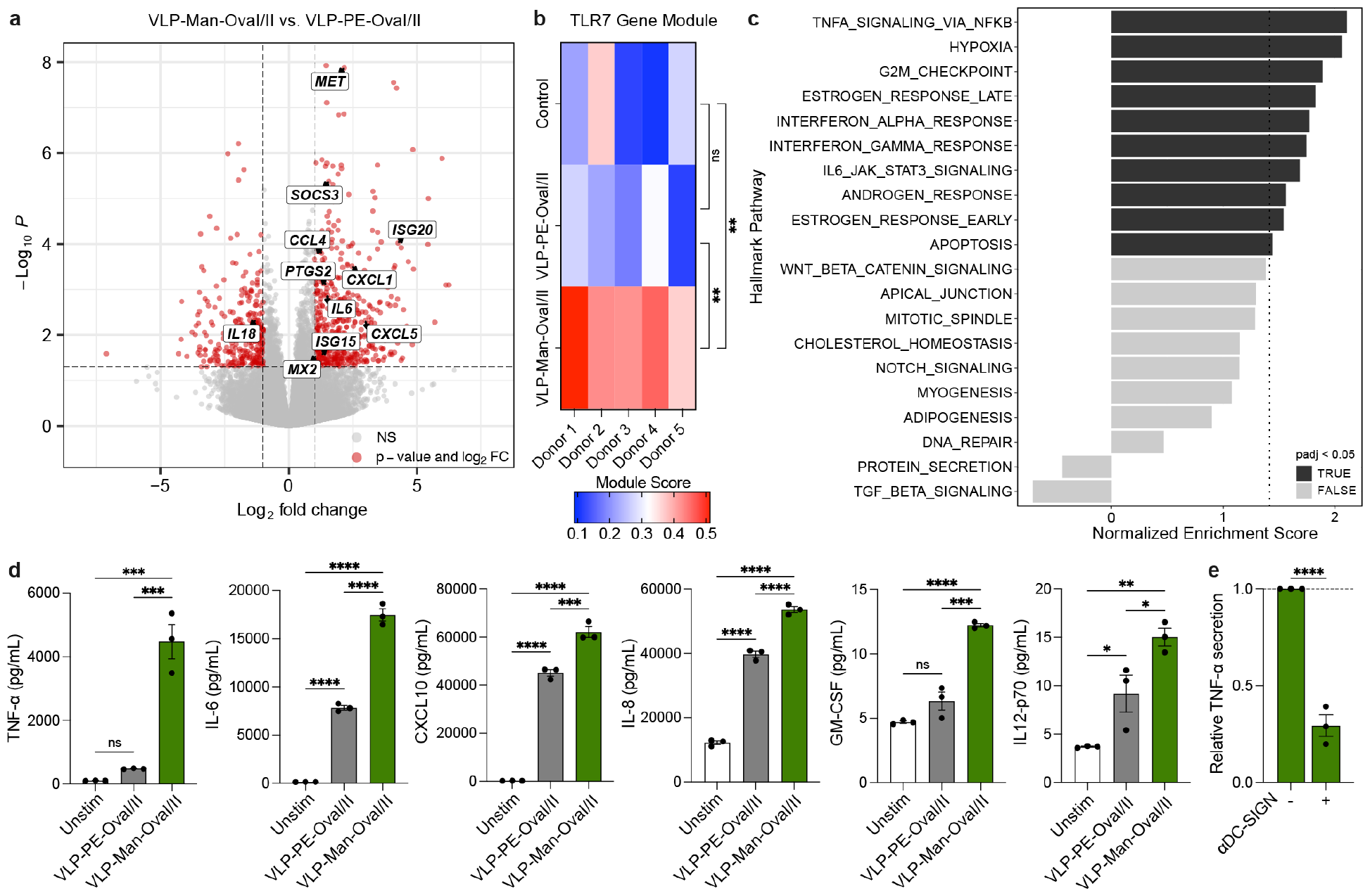
VLP-Man-OvaI/II induces type 1-associated responses. **a**, Altered gene expression profiles between VLP-Man-OvaI/II- and VLP-PE-OvaI/II-treated moDCs. moDCs were treated with 8 nM VLPs for 6 h. **b**, TLR7 downstream gene signature module score for unstimulated, VLP-PE-OvaI/II-, and VLP-Man-OvaI/II-treated moDCs. **c**, GSEA analysis of upregulated and downregulated pathways in DCs treated with VLP-Man-OvaI/II and VLP-PE-OvaI/II. **d**, The moDCs were treated with VLPs for 32 h and supernatants were collected for cytokine analysis. **e**, The moDCs were pre-treated with blocking antibodies against DC-SIGN for 20 min followed by stimulation with VLP-PE-OvaI/II or VLP-Man-OvaI/II for 32 h, and TNF-α secretion was measured. Data represent mean □ s.e.m. from a representative experiment (n=3). Statistical analysis was performed by one-way ANOVA with Bonferroni’s multiple comparisons test. *P<0.1, **P<0.01, ***P<0.001, ****P<0.0001. For **d** and **e**, experiments were repeated at least twice with similar results.

To assess downstream cytokine secretion, moDCs were incubated with VLPs for 32 h and the supernatant was collected for analysis. The moDCs exposed to VLP-Man-OvaI/II secreted significantly higher levels of type 1, pro-inflammatory cytokines, including TNF-α, IL-6, CXCL10, IL-8, and IFNλ1, compared to those treated with VLP-PE-OvaI/II (Fig. 3d). Notably, moDCs pre-treated with blocking antibodies against DC-SIGN showed a significant reduction in TNF-α production following VLP-Man-OvaI/II addition (Fig. 3e). This finding shows that engagement of DC-SIGN and TLR7 by VLP-Man-OvaI/II triggers a signaling cascade leading to type 1, pro-inflammatory cytokine secretion.

### Mannosylated VLPs are taken up preferentially by DCs *in vivo*

Motivated by the efficient uptake of mannosylated VLPs in moDCs, we examined the uptake of VLPs by DCs *in vivo*. Trafficking of vaccines to the lymph nodes (LNs) is a key step for efficient uptake by DCs and antigen presentation to T cells. Particles on the order of 20-200 nm have been shown to drain directly to LNs within hours of subcutaneous (s.c.) administration.^44^ The size of PP7 VLPs (∼50 nm diameter) is therefore ideal for entering the lymphatic system, rendering them capable of immune priming even when administered at sites distal from both the lymph vessels and the tumor.^44^ To assess the trafficking of the VLPs, C57BL/6J mice were immunized s.c. with AF647-labeled VLPs, and fluorescence intensity in the isolated organs was quantified. We observed selective accumulation of VLPs in the immune cell-rich LNs within 4 hours and clearance through the liver over the course of 72 hours (Supplementary Fig. 4a,b). We also inoculated mice with B16F10-OVA cells and allowed tumors to grow to ∼200 mm^3^ followed by s.c. administration of VLPs and measurement of VLP fluorescence intensity within the tumor over time. No localization of the VLPs within the tumor was observed, suggesting that there is no direct VLP activity within the tumor. (Supplementary Fig. 4c). Active targeting with mannoside ligands did not significantly alter VLP distribution to measured organs at 24 h (Supplementary Fig. 4d). Instead, mannosylation of VLPs enhanced their uptake by LN-resident DCs. VLP-Man-OvaI/II uptake was significantly higher than VLP-PE-OvaI/II uptake by DCs in the LNs after 24 h (Supplementary Fig. 4e,f). We hypothesized that this increased VLP-Man-OvaI/II uptake in combination with greater DC activation would facilitate more robust anti-tumor immunity.

### Mannosylated VLPs induce antigen-specific Th1 CD4^+^ and CD8^+^ T-cell responses *in vivo*

To evaluate tumor antigen-specific cellular immune responses, we examined the immunomodulatory effects of the VLPs in an immune-competent and syngeneic mouse transplant using the aggressive B16F10-OVA murine melanoma model. This solid tumor model expresses ovalbumin antigens and is non-responsive to anti-PD-1 treatment. We first investigated the ability of VLP-Man-OvaI/II to induce tumor antigen-specific CD4^+^ and CD8^+^ T cells that mediate tumor growth inhibition. A dinucleotide agonist for STING, 2′3′ cyclic GMP-AMP (cGAMP), was used as an adjuvant to boost T cell activation,^46, 47^ and the immune checkpoint inhibitor anti-PD-1 was used so that cytotoxic capacity of T cells could be directly measured. Six-week-old C57BL/6J mice were inoculated with B16F10-OVA cells, followed by three immunizations (at 6-day intervals) with VLPs, 2’3’-cGAMP, and anti-PD-1 (Fig. 4a,b). On day 21 (6 days following the last dose), spleens and tumors were isolated for immune cell analysis.

**Fig. 4.**
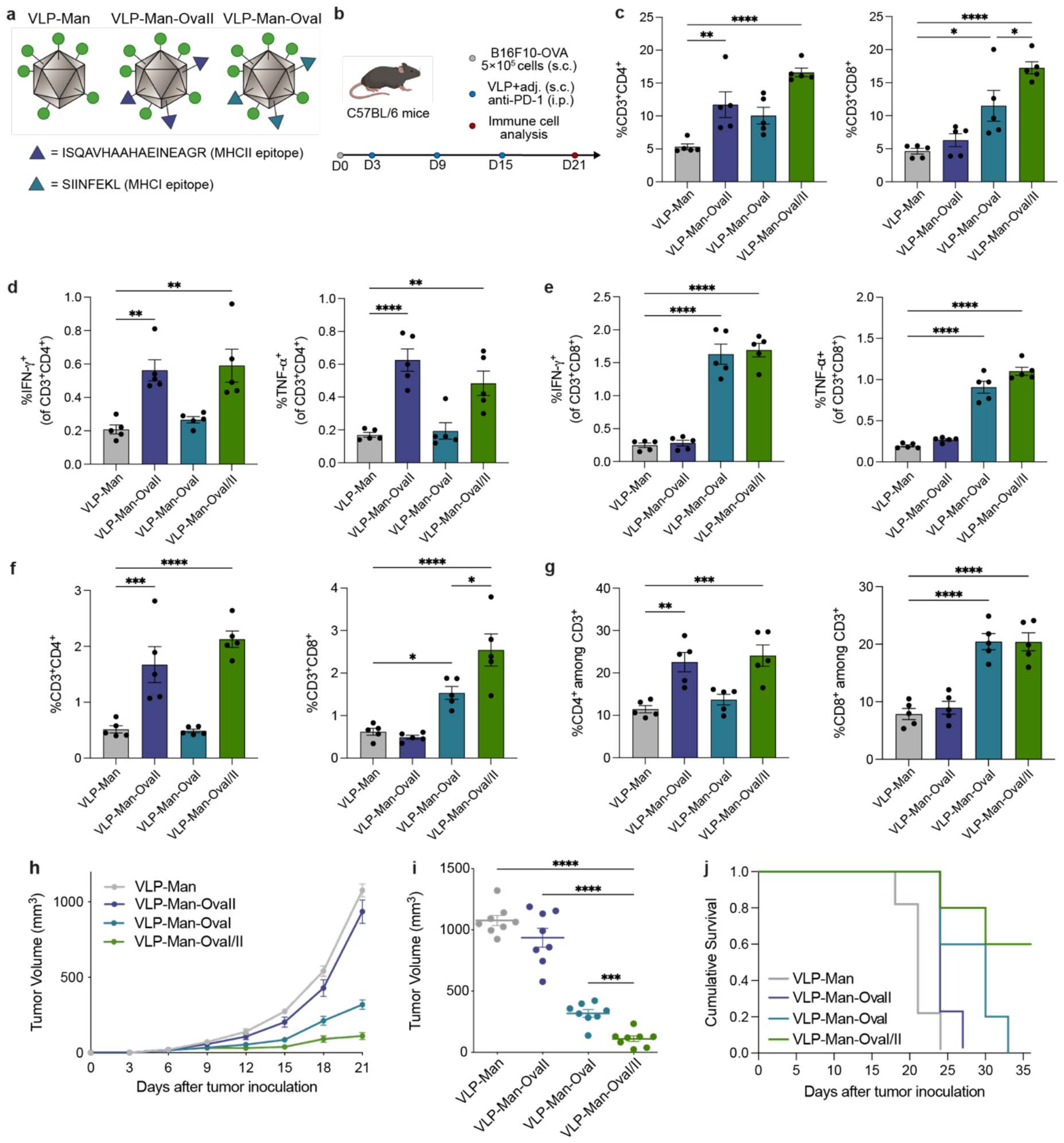
Mannosylated VLP conjugation with tumor antigens induces specific Th1 CD4^+^ and CD8^+^ T cells. **a**, Schematic representation of VLPs with Ova antigens. **b**, C57/BL6 mice were inoculated with 5×10^5^ B16F10-OVA cells (s.c.) on day 0 and treated with VLPs and 2’3’-cGAMP (s.c.) and anti-PD-1 intraperitoneally (i.p.) on days 3, 9, and 15. **c**, Percentages of CD4^+^ and CD8^+^ T cells in the spleens were analyzed on day 21. **d**, IFN-γ and TNF-α-secreting CD4^+^ T cells in the spleen were measured following restimulation with OVA(323-339). **e**, IFN-γ and TNF-α-secreting CD4^+^ T cells in the spleen were measured following restimulation with OVA(257-264). **f**, Percentages of CD4^+^ and CD8^+^ T cells in the tumors were analyzed on day 21. **g**, Percentages of CD4^+^ and CD8^+^ among CD3^+^ T cells within the tumor. **h**, Mean tumor growth curves. **i**, Tumor volumes on day 21. **j**, Survival curves. The difference between VLP-Man and VLP-Man-OvaI/II was significant (p < 0.01, Log-rank (Mantel– Cox) test). Data represent mean ± s.e.m. from a representative experiment (n=5 (**c**-**g**) and n=8 (**h**-**j**). Statistical analysis was performed by one-way ANOVA with Bonferroni’s multiple comparisons test or by log-rank (Mantel-Cox) test for the survival analysis. *P<0.1, **P<0.01, ***P<0.001, ****P<0.0001. Each experiment was repeated twice with similar results.

We first measured the ability of mannosylated VLPs conjugated with tumor antigens to induce tumor-antigen specific Th1 CD4^+^ and CD8^+^ T cells. To this end, we compared T cell induction by VLP-Man, VLP-Man-OvaII, VLP-Man-OvaI, and a combination of both VLPs (VLP-Man-OvaI/II) (Fig. 4a). Immunization with VLP-Man-OvaII and VLP-Man-OvaI resulted in increased percentages of CD4^+^ and CD8^+^ T cells, respectively, compared to immunization with VLP-Man only (Fig. 4c). Interestingly, immunization with the combination particles VLP-Man-OvaI/II led to a significantly higher percentage of CD8^+^ T cells in the spleen compared to VLP-Man-OvaI alone (Fig. 4c), in line with ability of Th1 CD4^+^ T cell help to augment CD8^+^ T cell proliferation.^48^ To assess T cell specificity against tumor antigens, we restimulated splenocytes with the antigens OVA(323-339) or OVA(257-264) and used intracellular cytokine staining to quantify the percentage of CD4^+^ and CD8^+^ T cells expressing IFN-γ and TNF-α, cytokines characteristic of type 1 polyfunctional T cells. Upon restimulation with OVA(323-339), a significant increase in the percentage of IFN-γ^+^CD4^+^ T cells and TNF-α^+^CD4^+^ T cells among spleen cells was detected in the VLP-Man-OvaII immunization group compared to those treated with VLP-Man alone (Fig. 4d). Notably, a significant increase in IFN-γ^+^CD8^+^ T cells (6.6-fold) and TNF-α^+^CD8^+^ T cells (4.7-fold) among splenocytes was detected in the Man-OvaI group compared to the Man group upon restimulation with OVA(257-264) (Fig. 4e). Both tumor-antigen specific CD4^+^ and CD8^+^ T cells were induced by immunization with VLP-Man-OvaI/II (Fig. 4d,e). Thus, simultaneous presentation of exogenous glycoligands with tumor antigens by VLPs enabled the differentiation of naïve T cells into tumor antigen-specific Th1 CD4^+^ and CD8^+^ T cells *in vivo*.

The extent of immune infiltrate within tumors is emerging as one of the critical predictors of cancer progression, with the presence of tumor-infiltrating lymphocytes (TILs) being correlated to patient survival.^49,50^ We therefore assessed T cell migration to the tumor site. Consistent with T cell induction in the spleen, we observed an upregulation of CD4^+^ and CD8^+^ T cells within the tumor corresponding to immunization with VLP-Man-OvaII, VLP-Man-OvaI, or VLP-Man-OvaI/II compared to immunization with Man alone (Fig. 4f,g). Additionally, we observed a significant increase in the percentage of CD8^+^ T cells in the tumors of mice immunized with VLP-Man-OvaI/II compared to Man-OvaI, once again suggesting Th1 CD4^+^ T cell help promoted CD8^+^ T cell proliferation (Fig. 4f).

Tumor growth was monitored over the course of 3 weeks to assess the cytotoxic capacity of these T cells. Immunization with VLP-Man-OvaII did not provide significant tumor growth inhibition compared to VLP-Man only, suggesting that an enhanced CD4^+^ T cell response alone is not sufficient for anti-tumor immunity (Fig. 4h,i). In contrast, VLP-Man-OvaII controls provided significant tumor growth inhibition over VLP-Man only, demonstrating that the resulting CD8^+^ T cells were able to achieve significant tumor cell killing (Fig. 4h,i). Finally, VLP-Man-OvaI/II significantly inhibited tumor growth and prolonged animal survival compared to all other cohorts (Fig. 4h-j), suggesting that holistic induction of Th1 CD4^+^ and CD8^+^ T cells provides enhanced immunity. Overall, we demonstrated tumor antigen-specific T cell responses that were directly measurable in the splenocytes and tumors of mice following vaccination with VLP-Man-OvaI/II, resulting in therapeutic efficacy in a murine melanoma model.

### Mannosylation amplifies anti-tumor efficacy of VLPs

The engagement of DC-SIGN by VLP-Man-OvaI/II induced differential DC activation and signaling profiles *in vitro* compared to the nonspecific PE particles. We next assessed if this superior DC activation translated to a more robust T cell response *in vivo* relative to an immunogenic presentation of the target tumor antigens alone. Thus, while administration of VLP-PE-OvaI/II slowed tumor growth, vaccination with VLP-Man-OvaI/II, displaying the DC-SIGN ligand, significantly inhibited tumor growth and extended survival in the B16F10-OVA model (Fig. 5a-d). Examination of the immune landscape on day 21 revealed that DC activation paralleled T cell generation and therapeutic efficacy. Immunization with VLP-Man-OvaI/II significantly increased percentages of both CD4^+^ and CD8^+^ T cells in the spleen compared to immunization with VLP-PE-OvaI/II (Fig. 5e). Not only did the mannosylated particles recruit more T cells to the spleen, but they also induced a significantly higher percentage of tumor antigen-specific T cells compared to the control particles. Specifically, IFN-γ and TNF-α secretion upon restimulation with OVA(323-339) or OVA(257-264) were significantly higher for mice immunized with VLP-Man-OvaI/II (Fig. 5f,g). Compared to the VLP-PE-OvaI/II control, we detected a significant increase in the percentage of CD4^+^ and CD8^+^ T cells among all tumor cells and among CD3^+^ T cells in the tumors of VLP-Man-OvaI/II immunized mice (Fig. 5h,i). Consistent with our *in vitro* results, these data suggest that the mannosylated VLPs drive greater DC activation, antigen presentation, and more potently activating immune signaling to enhance T cell responses.

**Fig. 5.**
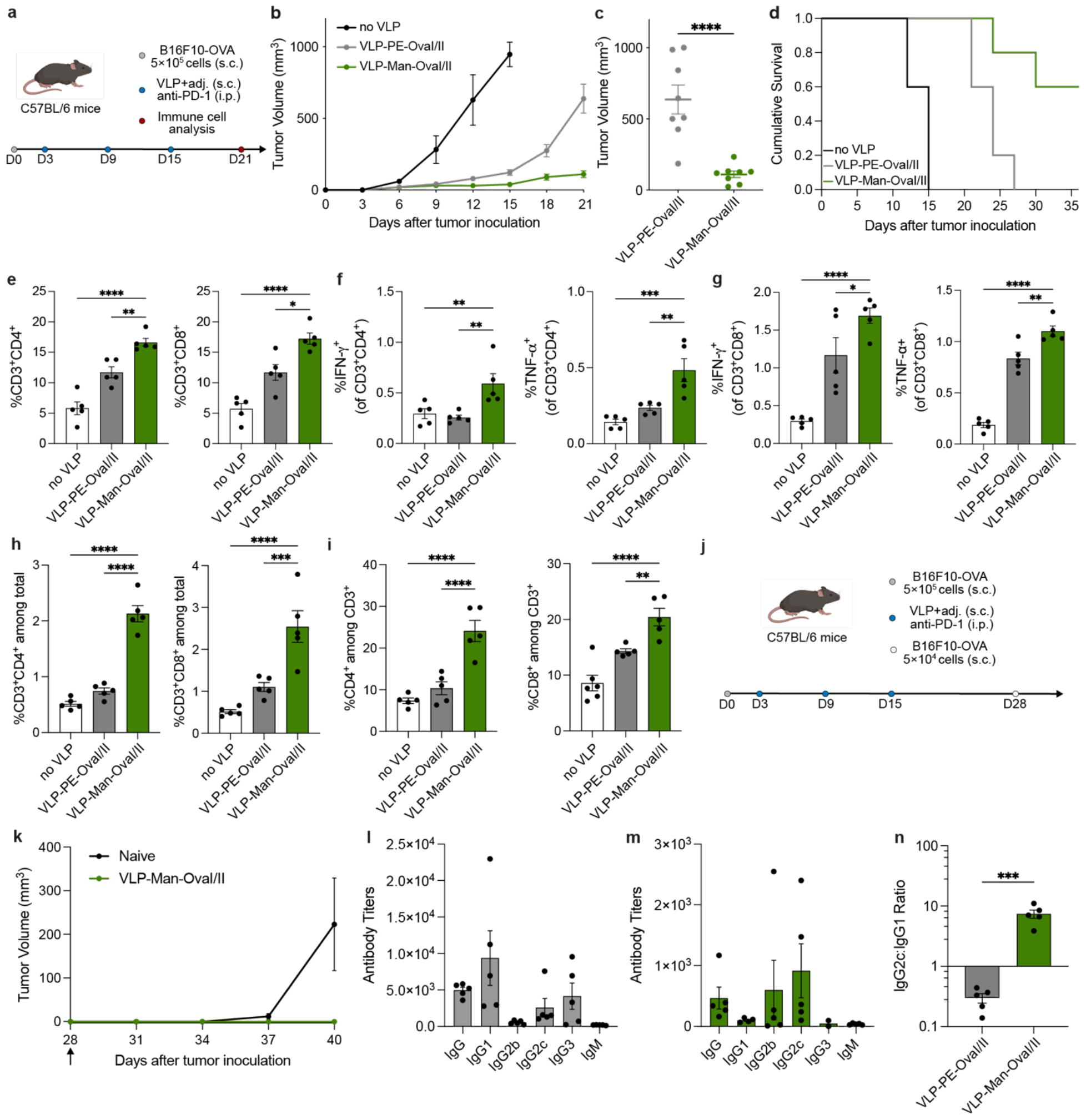
VLP-Man-OvaI/II induces tumor antigen-specific T cells and exerts anti-tumor response. **a**, C57/BL6 mice were inoculated with 5×10^5^ B16F10-OVA cells (s.c.) on day 0 and treated with VLPs and 2’3’-cGAMP (s.c.) and anti-PD-1 intraperitoneally (i.p.) on days 3, 9, and 15. **b**, Mean tumor growth curves. **c**, Tumor volumes on day 21. **d**, Survival curves. The difference between VLP-PE-OvaI/II and VLP-Man-OvaI/II was significant (p < 0.1, Log-rank (Mantel–Cox) test). **e**, Percentages of CD4^+^ and CD8^+^ T cells in the spleens were analyzed on day 21. **f**, IFN-γ and TNF-α-secreting CD4^+^ T cells in the spleen were measured following restimulation with OVA(323-339). **g**, IFN-γ and TNF-α-secreting CD8^+^ T cells in the spleen were measured following restimulation with OVA(257-264). **h**, Percentages of CD4^+^ and CD8^+^ T cells in the tumors were analyzed on day 21. **i**, Percentages of CD4^+^ and CD8^+^ among CD3^+^ T cells within the tumor. **j**, Surviving mice were re-challenged with 5×10^4^ B16F10-OVA cells on day 28. **k**, Tumor growth curves. **l**,**m**, OVA(323-339)-specific antibody profiles were analyzed by ELISA measurements from serum of mice immunized with VLP-PE-OVAI/II (**l**) or VLP-Man-OvaI/II (**m**). **n**, IgG2c:IgG1 antibody ratio for mice immunized with VLPs. Line represents IgG2c:IgG1 ratio of 1. Data represent mean +/-s.e.m. from a representative experiment (n=8 (**b-d**) and n=5 (**e-i**) and n=4 (**j-k**) and n=5 (**l-n**)). Statistical analysis was performed by one-way ANOVA with Bonferroni’s multiple comparisons test or by log-rank (Mantel-Cox) test for the survival analysis. *P<0.1, **P<0.01, ***P<0.001, ****P<0.0001. Each experiment was repeated twice with similar results.

As melanoma is a highly metastatic and recurrent disease, robust protection against melanoma recurrence is needed.^51^ To test our VLP strategy on this front, surviving mice from the VLP-Man-OvaI/II cohort were re-challenged with B16F10-Ova on Day 28 (Fig. 5j). While naïve mice rapidly developed tumors, the VLP-Man-OvaI/II cohort was protected against tumor re-challenge (Fig. 5k). The immune protection observed following therapeutic vaccine administration suggests the generation of long-term memory T cells that provide prophylactic immunity and combat disease recurrence.

We also investigated humoral responses by analyzing antibody profiles of mice immunized with VLPs. OVA(323-339)-specific antibodies were evaluated on day 21. Serum ELISA analyses showed that VLP-Man-OvaI/II-immunized mice produced lower OVA(323-339)-specific IgG antibody titers than mice immunized with VLP-PE-OvaI/II (Fig. 5l-m). This decrease in antibody production is consistent with the expectation that VLP-Man-OvaI/II biases the immune response towards type 1 cellular immunity and away from type 2 humoral immunity involving antibody production. Furthermore, VLP-PE-OvaI/II and VLP-Man-OvaI/II immunization induced distinct antibody profiles—VLP-PE-OvaI/II primarily induced IgG1-type antibodies while VLP-Man-OvaI/II primarily induced IgG2b/c antibodies (Fig. 5l-m). We evaluated IgG2c/IgG1 ratios, as higher ratios are indicative of a Th1-type immune response. Mice immunized with VLP-Man-OvaI/II displayed an antibody subclass IgG2c/IgG1 ratio greater than 1, while mice immunized with VLP-PE-OvaI/II displayed a ratio less than 1 (Fig. 5n). These data are consistent with the increased T cell responses observed with immunization by VLP-Man-OvaI/II, confirming a shift from humoral immunity to cellular immunity via engagement of DC-SIGN. Collectively, these results demonstrate that immunization with VLP-Man-OvaI/II generates robust, tumor antigen-specific Th1 CD4^+^ and CD8^+^ T cells capable of tumor infiltration and yielding potent anti-tumor responses.

### VLP-Man-OvaI/II serves as a self-adjuvanting vaccine

Having demonstrated that VLP-Man-OvaI/II vaccination induces tumor-antigen specific Th1 CD4^+^ and CD8^+^ T cells, we sought to assess the potential benefits of simultaneous STING activation with the adjuvant cGAMP. Mice inoculated with B16F10-OVA tumor cells were immunized with VLP-Man-OvaI/II and anti-PD-1 with and without the adjuvant 2’3’-cGAMP (Fig. 6a). No significant difference in tumor growth or animal survival was observed (Fig. 6b-d). Immune cell analysis showed a decrease in the percentage of antigen-specific CD4^+^ T cells and no significant difference in the percentage of antigen-specific CD8^+^ T cells (Fig. 6e-f). The same trends were observed in the tumor, with a decrease in the percentage of tumor-infiltrating CD4^+^ T cells and no significant decrease in the percentage of tumor-infiltrating CD8^+^ T cells (Fig. 6g). These data indicate that VLP-Man-OvaI/II can serve as a self-adjuvanting cancer vaccine due to the immunostimulatory signaling achieved via combinatorial engagement of TLR7 and DC-SIGN.

**Fig. 6.**
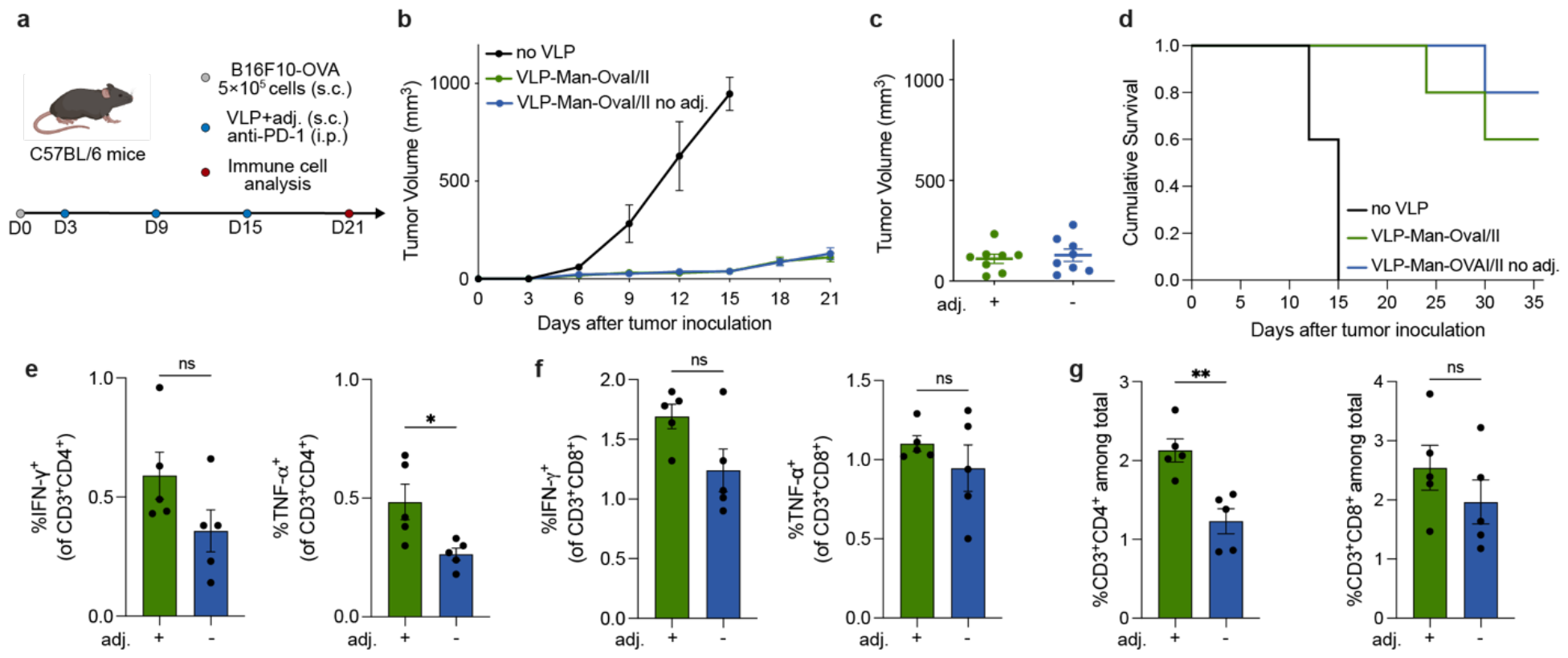
VLP-Man-OvaI/II has self-adjuvanting capacity. **a**, C57/BL6 mice were inoculated with 5×10^5^ B16F10-OVA cells (s.c.) on day 0 and treated with VLPs and 2’3’-cGAMP (s.c.) and anti-PD-1 intraperitoneally (i.p.) on days 3, 9, and 15. **b**, Mean tumor growth curves. **c**, Tumor volumes on day 21. **d**, Survival curves. Differences between VLP-Man-OvaI/II and VLP-Man-OvaI/II no adj. were not significant. **e**, IFN-γ and TNF-α-secreting CD4^+^ T cells in the spleen were measured following restimulation with OVA(323-339). **f**, IFN-γ and TNF-α-secreting CD8^+^ T cells in the spleen were measured following restimulation with OVA(257-264). **g**, Percentages of CD4^+^ and CD8^+^ T cells in the tumors were analyzed on day 21. Data represent mean +/-s.e.m. from a representative experiment (n=8 (**b-d**) and n=5 (**e-g**)). Statistical analysis was performed by one-way ANOVA with Bonferroni’s multiple comparisons test or by log-rank (Mantel-Cox) test for the survival analysis. *P<0.1, **P<0.01, ***P<0.001, ****P<0.0001.

## Discussion

Effective cancer vaccines require mobilization of type 1 cellular immunity, a process that necessitates DC recognition of endogenous tumor antigens as pathogenic.^52^ Currently FDA-approved cancer vaccines involve adoptive transfer of *ex vivo* loaded DCs;^53^ however, these individualized procedures are complex, lengthy, and expensive. Moreover, significant limitations of this approach, including poor immune cell persistence and low tumor infiltration, result in limited tumor control.^54^ To overcome these challenges, there is a critical need for strategies that facilitate *in vivo* priming of DCs and T cells against tumor antigens.^2^ Achieving the desired T cell polarization *in vivo* requires localization to the LNs, activation of distinct DC signaling pathways, and secretion of pro-inflammatory cytokines.^55^ While antibody conjugates efficiently target DC-specific lectins for tumor antigen delivery, they do not achieve sufficient lectin clustering to induce cellular immunity.^56^ On the other hand, TLR agonists have demonstrated promise in promoting DC activation and cellular immunity, but their application has been limited by poor efficacy and systemic toxicity.^20, 24^ Enhancing delivery to APCs via particulate formulations of TLR agonists can increase efficacy while minimizing off-target effects.^57, 58^ A strategy that combines lectin and TLR signaling holds promise to facilitate *in vivo* priming of DCs and T cells against tumor antigens.

Mannose functionalization has previously been utilized to increase uptake of TLR agonists via nonselective targeting of lectins on APCs with mixed results.^25-29^ In addition to utilizing lectins to facilitate DC uptake and minimize off-target activity, we sought to methodically leverage defined pathogen-mimetic glycans to selectively engage immune activating lectins, initiate lectin signaling, and amplify TLR activation. Indeed, our previous studies demonstrated that the chemical structure of mannose ligands has profound effects on downstream immunity. Specifically, we observed an augmentation of Th1 responses only with an aryl mannoside ligand that binds DC-SIGN in a pH-independent manner, while pH-dependent mannose ligands only increased antigen uptake without enhancing immune responses.^31^ We therefore hypothesized that leveraging lectin-driven immune signaling could amplify TLR activation to induce robust anti-tumor cellular immunity.

Our vaccine platform is uniquely designed to emulate exogenous pathogen signaling to reshape the immune response against endogenous tumor antigens. To achieve this pathogen mimicry, we engineered DC-SIGN-targeted VLPs that package ssRNA as masked TLR ligands. The self-assembled PP7 VLP was functionalized with exogenous mannose ligands and model tumor antigens to yield the vaccine candidate VLP-Man-OvaI/II. The use of a synthetically accessible aryl mannoside derivative that binds the lectin DC-SIGN in a pH-independent manner facilitated efficient DC-SIGN engagement and signaling, without requiring isolation of native and otherwise prohibitively complex carbohydrates.^31^ Attachment via powerful azide-alkyne bioconjugation chemistry allows the required high density of ligand presentation to be achieved in a manner compatible with the co-display of antigenic peptides. The VLPs traffic to the immune cell-rich LNs following subcutaneous administration, enabling induction of tumor-specific immunity without requiring intratumoral administration. This is advantageous to overcome limited tumor accessibility, the tolerogenic tumor environment, acceptability of delay before tumor excision, and tumor metastasis.^59^

DC-SIGN-mediated endocytosis facilitated highly efficient uptake of VLP-Man-OvaI/II and TLR7 recognition of encapsulated ssRNAs. RNA-Seq analysis revealed that VLP-Man-OvaI/II upregulated TLR7-associated gene expression compared to identical particles that do not engage DC-SIGN. Additionally, VLP-Man-OvaI/II induced higher levels of DC activation and pro-inflammatory cytokine secretion compared to the control VLP-PE-OvaI/II, demonstrating the benefit of simultaneously delivering ligands for DC-SIGN and TLR7. Efficient presentation of tumor antigens on activated DCs induced tumor antigen-specific Th1 CD4^+^ and CD8^+^ T cells capable of tumor infiltration. We demonstrated that Th1 CD4^+^ T cell help supported CD8^+^ T cell proliferation, survival, and effector functions. Immunization with VLP-Man-OvaI/II significantly inhibited tumor growth and prolonged survival compared to VLP-PE-OvaI/II in a challenging murine melanoma model. Generation of tumor antigen-specific T cells *in vivo* established immune memory, providing protection against tumor re-challenge. This capability to prevent tumor recurrence following treatment adds significant value to our therapeutic cancer vaccine. Finally, we demonstrated the capacity of VLP-Man-OvaI/II to serve as a self-adjuvanting cancer vaccine, preventing the need for additional adjuvants that could overstimulate the immune system.

Our novel platform delivers tumor antigens in a pathogen costume, mimicking key features of extracellular pathogens to drive immunity against cancer cells. We harnessed pathogenic glycosylation to boost immune responses against otherwise tolerated tumor antigens, contrary to the glycan-lectin interactions commonly exploited by cancer to evade immunity. Our vaccine system mirrors the immunostimulatory signaling of foreign pathogens via simultaneous delivery of DC-SIGN and TLR7 agonists to trigger immune activation against tumor antigens.

These findings constitute a significant conceptual advance in cancer vaccine development by leveraging specific lectin signaling to drive TLR activation and anti-tumor immunity. The strategy we described shows promise as a viable alternative to autologous cell immunotherapies, possessing the ability to induce robust tumor antigen-specific T cells capable of solid tumor infiltration. The straightforward conjugation strategy used for VLP functionalization enables the installation of multiple human tumor neoantigens, facilitating potential translation of this platform to a clinical setting. This work has the potential to broaden the scope of cancer immunotherapies by presenting a modular and synthetically accessible platform capable of directing type 1 cellular immunity. Finally, we envisage that this concept could be applied to vaccine development for diverse intracellular infectious diseases that have eluded vaccine realization.

## Acknowledgements

This research was supported by the National Institutes of Health (R01A1055258 to L.L.K, R01CA247484 to M.G.F., 2R01CA220468-06A1 to J.A.J., and F99CA264404 to S.H.B.), the Centers for Disease Control and Prevention (75D30121P12509 to M.G.F.), the Marie Skłodowska-Curie Individual Fellowship (MSCA-IF-GF 886709 to A.G.), and the National Science Foundation (NSF GRFP fellowship for V.L.). We thank the Koch Institute’s Robert A. Swanson (1969) Biotechnology Center, specifically the Flow Cytometry Core and the Integrated Genomics and Bioinformatics Core, supported in part by the Koch Institute Support Grant P30-CA014051 from the National Cancer Institute. We also thank the MIT Department of Chemistry Instrumentation Facility, the MIT Division of Comparative Medicine, and the Imaging Core of the Ragon Institute of MGH, MIT, and Harvard. We thank T. Diefenbach for assistance with microscopy. Figure 1a. was created with BioRender.com.

## Competing Interests

A.K.S. reports compensation for consulting and/or Scientific Advisory Board (SAB) membership from Merck, Honeycomb Biotechnologies, Cellarity, Repertoire Immune Medicines, Hovione, Third Rock Ventures, Ochre Bio, FL82, Empress Therapeutics, Relation Therapeutics, Senda Biosciences, IntrECate biotherapeutics, Santa Ana Bio, and Dahlia Biosciences unrelated to this work. D.J.I. reports compensation for consulting and/or SAB membership from Elicio Therapeutics, Ankyra Therapeutics, Strand Therapeutics, Window Therapeutics, Venn Therapeutics, Alloy Therapeutics, Livzon Pharmaceuticals, SQZ Biotechnologies, Jupiter Therapeutics, Parallel Bio, Surge Therapeutics, Senda Biosciences, Gensaic Therapeutics, and Third Rock Ventures unrelated to this research. J.A.J. is a cofounder and shareholder of Window Therapeutics unrelated to this research. L.L.K. reports compensation for consulting and/or SAB membership from Exo Therapeutics, the ONO Pharmaceutical Foundation, and Coca Cola unrelated to this research. V.L., R.H., M.M.A., L.L.K., and M.G.F. are inventors on relevant patent applications held by the Massachusetts Institute of Technology and Georgia Institute of Technology. The remaining authors declare that they have no competing interests.

