## Supplementary Information for "Glycan-costumed virus-like particles promote type 1 anti-tumor immunity"

**This PDF file includes:**
Materials and Methods
Scheme S1
Table S1 to S2
Figs. S1 to S4

### Materials and Methods

*General information.* All reagents and solvents were purchased from Sigma Aldrich and used without further purification unless otherwise noted. Dry solvents were obtained from a solvent purification system (Pure Process Technologies). Analytical thin layer chromatography (TLC) was used to monitor reactions and was performed on EMD Millipore TM TLC silica gel 60 F254 (glass-backed). Compounds were visualized with ultraviolet light (254 nm) and/or charring with *p*-anisaldehyde. Flash chromatography was performed on SiliCycle ® SiliaFlash ® P60 silica gel and Biotage ® Selekt using Biotage Sfär silica cartridges. High-performance liquid chromatography (HPLC) was performed on a 1260 Infinity II Preparative LC System (Agilent Technologies) equipped with a XSelect Peptide CSH C18 OBD Prep Column (130 Å pore size, 5 □m particle size, 19 mm × 250 mm of width × length) (Waters Corporation, Milford, MA).

$^1H$ and $^{13}C$ nuclear magnetic resonance (NMR) spectra were acquired at the MIT Department of Chemistry Instrumentation Facility on a Bruker Avance Neo 600 MHz spectrometer. Chemical shifts were reported relative to residual solvent peaks in parts per million ($CHCl_3$: $^1H$ δ 7.26, $^{13}C$ δ 77.0; $CH_3OH$: $^1H$ δ 3.31, $^{13}C$ δ 49.0). Peak multiplicity is reported as singlet (s), doublet (d), multiplet (m), doublet of doublet (dd), etc. All spectra are reported in the supplemental information. Electrospray ionization mass spectra (ESI-MS) were obtained on an Agilent 6125B mass spectrometer attached to an Agilent 1260 Infinity LC System**.**

### Synthetic procedures and characterization data

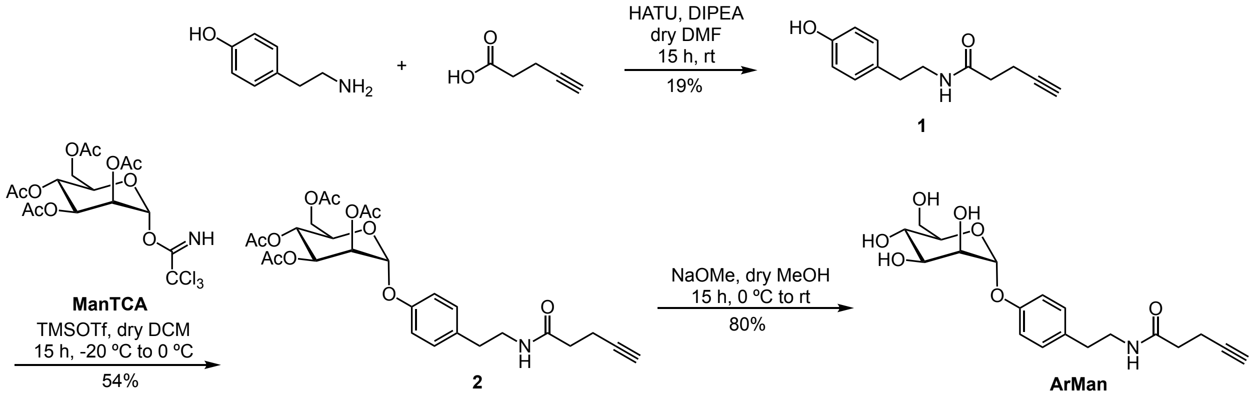


**Scheme S1.** Synthesis of aryl mannoside ligand.

(Scheme 1) *N-(4-hydroxyphenethyl)pent-4-ynamide,* **1**. Pent-4-ynoic acid (0.284 g, 2.90 mmol) was dissolved in dry, deoxygenated DMF (5 mL) in a flame-dried flask. 1-[Bis(dimethylamino)methylene]-1*H*-1,2,3-triazolo[4,5-*b*]pyridinium 3-oxide hexafluorophosphate (1.102 g, 2.90 mmol) and *N*,*N*-diisopropylethylamine (0.637 mL, 2.90 mmol) were added and stirred for 5 min. Tyramine (0.400 mg, 2.92 mmol) was added, and the solution was stirred for 15 h at rt. The reaction mixture was concentrated under reduced pressure, and the product was purified by normal phase flash chromatography eluting with an EtOAc/hexanes gradient (0:1 to 1:0) to afford the alkyne tyramide intermediate (**1**, 0.120 g, 0.551 mmol, 19%). $^1H$ NMR (600 MHz, $CDCl_3$) δ 7.04 (d, $J$ = 8.4 Hz, 2H), 6.79 (d, $J$ = 8.4 Hz, 1H), 5.70 (br s, 1H), 3.51 (q, $J$ = 6.8 Hz, 2H), 2.75 (t, $J$ = 6.9 Hz, 2H), 2.50 (td, $J$ = 7.2, 2.6 Hz,

2H), 2.36 (t, $J$ = 7.2 Hz, 2H), 1.96 (t, $J$ = 2.6 Hz, 1H). $^{13}$C NMR (150 MHz, $CDCl_3$) δ 171.4, 154.8, 130.4, 130.0, 115.7, 83.0, 70.0, 41.1, 35.5, 34.8, 15.0. Calculated m/z for $C_{13}H_{15}NO_2$: $[M+H]^+$, 218.1; found 218.1. The recorded spectra of **1** matched those previously reported in literature.[1]

(Scheme 1) *(2R,3R,4S,5S,6R)-2-(acetoxymethyl)-6-(4-(2-(pent-4-ynamido)ethyl)phenoxy) tetrahydro-2H-pyran-3,4,5-triyl triacetate*, **2**. 2,3,4,6-Tetra-O-acetyl-α-D-mannopyranosyl trichloroacetimidate (**ManTCA**) was synthesized as previously reported.[2] **ManTCA** (0.150 g, 0.304 mmol) and compound **1** (0.074 g, 0.340 mmol) were dissolved in dry $CH_2Cl_2$ (5 mL) and stirred with 4 Å molecular sieves at rt for 1 h. The suspension was cooled to -20 °C, and trimethylsilyl trifluoromethanesulfonate (44 µL, 0.243 mmol) was added. The reaction mixture was stirred for 15 h at 0 °C and then quenched with triethylamine to neutral pH. The mixture was diluted with $CH_2Cl_2$, and molecular sieves were removed by filtration. The organic phase was washed with water, dried over $MgSO_4$, filtered, and concentrated under reduced pressure. The crude oil was purified by column chromatography (3:7 EtOAc:hexanes) to afford compound **2** (0.090 g, 0.164 mmol, 54%) as a colorless oil. $^1$H NMR (600 MHz, $CDCl_3$) δ 7.13 (d, $J$ = 8.6 Hz, 2H), 7.02 (d, $J$ = 8.6 Hz, 2H), 5.62 (br t, $J$ = 5.8 Hz, 1H), 5.55 (dd, $J$ = 10.1, 3.5 Hz, 1H), 5.48 (d, $J$ = 1.9 Hz, 1H), 5.42 (dd, $J$ = 3.6, 1.9 Hz, 1H), 5.37 (t, $J$ = 10.0 Hz, 1H), 4.28 (dd, $J$ = 12.0, 5.0 Hz, 1H), 4.14 – 4.03 (overlapping signals, 2H), 3.50 (q, $J$ = 6.7 Hz, 2H), 2.78 (t, $J$ = 7.0 Hz, 2H), 2.50 (td, $J$ = 7.2, 2.7 Hz, 2H), 2.35 (t, $J$ = 7.2 Hz, 2H), 2.19 (s, 3H), 2.05 (s, 3H), 2.04 (s, 3H), 2.03 (s, 3H), 1.96 (t, $J$ = 2.6 Hz, 1H). $^{13}$C NMR (150 MHz, $CDCl_3$) δ 171.0, 170.7, 170.2, 170.1, 169.9, 154.6, 133.6, 130.0, 116.9, 96.1, 83.1, 69.6, 69.5, 69.2, 69.0, 66.1, 62.3, 40.9, 35.5, 35.0, 21.0, 20.9, 20.8, 20.8, 15.0. Calculated m/z for $C_{27}H_{23}NO_{11}$: $[M+H]^+$, 548.2; found 548.2.

(Scheme 1) *N-(4-(((2R,3S,4S,5S,6R)-3,4,5-trihydroxy-6-(hydroxymethyl)tetra-hydro-2H-pyran-2-yl)oxy)phenethyl)pent-4-ynamide*, **ArMan**. Compound **2** (0.090 g, 0.164 mmol) was dissolved in dry MeOH and freshly prepared sodium methoxide was added to reach a pH of 9 at 0 °C. The solution was allowed to stir for 15 h at rt. The reaction was neutralized with acetic acid, concentrated by rotary evaporation, then dissolved in water and lyophilized. The product was purified by HPLC using a gradient of 5-70% v/v ACN in $H_2O$ containing TFA (0.1% v/v) over 43 min. **ArMan** has a retention time of 11 minutes and fraction purity was assessed by LC-MS. Pure fractions were pooled and lyophilized to yield **ArMan** as the pure α anomer (0.049 g, 0.131 mmol, 80%). $^1$H NMR (600 MHz, MeOD) δ 7.15 (d, $J$ = 8.6 Hz, 2H), 7.04 (d, $J$ = 8.6 Hz, 2H), 5.43 (d, $J$ = 1.9 Hz, 1H), 3.98 (dd, $J$ = 3.4, 1.9 Hz, 1H), 3.89 (dd, $J$ = 9.5, 3.4 Hz, 1H), 3.78 – 3.68 (overlapping signals, 3H), 3.60 (ddd, $J$ = 9.8, 5.2, 2.6 Hz, 1H), 3.37 (t, $J$ = 7.4 Hz, 2H), 2.74 (t, $J$ = 7.3 Hz, 2H), 2.44 (td, $J$ = 7.5, 2.4 Hz, 2H), 2.34 (t, $J$ = 7.0 Hz, 2H), 2.26 (t, $J$ = 2.7 Hz, 1H). $^{13}$C NMR (151 MHz, MeOD) δ 174.0, 156.6, 134.4, 130.9, 117.9, 100.3, 83.5, 75.3, 72.4, 72.1, 70.3, 68.3, 62.7, 42.2, 36.0, 35.7, 15.7. Calculated m/z for $C_{19}H_{25}NO_7$: $[M+H]^+$, 380.1; found 380.1. The recorded spectra of **ArMan** matched those previously reported in literature.[3]

**Expression and purification of VLPs**

Dimeric PP7-PP7 VLPs were expressed in BL21(DE3) *Escherichia coli* (Biogen) and isolated as previously described.[4] A single isolated colony from a plate of transformed cells was grown overnight in 2xYT media supplemented with kanamycin (50 µg/mL). These starter cultures were used to inoculate larger cultures; cultures were grown to OD600 ~0.8, and protein expression was induced by addition of isopropyl β-D-1-thiogalactopyranoside (IPTG) to a final concentration of 1 mM. Following induction, cells were incubated at 37 °C for 4 h and harvested by centrifugation (6,000 rpm, 10 min). Cell pellets were used immediately or stored at −80 °C until processed.

Cell pellets were resuspended in 0.1 M potassium phosphate buffer, pH 7.4 (30-40 mL buffer per pellet from 250 mL cell culture). Resuspended cells were lysed by probe sonication (10 min on total, 50-60 W, 5 s on, 5 s off) in an ice bath, and clarified by centrifugation (14,000 rpm, 10 m). Protein in the resulting supernatant was precipitated by addition of 0.27 g/mL ammonium sulfate, with gentle rocking for 2 h at 4 °C, followed by centrifugation (14,000 rpm, 10 m). The resulting protein pellet was gently resuspended in 0.1 M potassium phosphate buffer, pH 7.4. Lipids and membrane proteins carried through precipitation were removed by organic extraction using one volume of 1:1 n-butanol/chloroform. The VLP-containing aqueous layer was removed following centrifugation (14,000 rpm, 10 min), and particles were further purified by 10–40% sucrose gradient ultracentrifugation (28,000 rpm, 4 h). Visible-particle-containing bands were extracted, and particles were isolated by ultracentrifugation (68,000 rpm, 2 h). VLP pellets were decanted, resuspended in 0.1 M potassium phosphate buffer, and sterilized by 0.2 μm PTFE syringe filters.

**Characterization of PP7-PP7 VLPs**

Protein concentration was determined by a Bradford assay (Pierce, Coomassie Plus) against BSA standards. Particles were characterized by FPLC using a Superose 6 size exclusion column (Cytiva) to determine particle purity, and by dynamic light scattering using a Dynapro plate reader (Wyatt) to determine hydrodynamic radius. Where reported, molar values for protein concentration refer to the concentration of intact particles, each composed of 120 self-assembling coat protein subunits.

**LC-MS analysis**

The extent of particle modification was assessed by LC-ESI-TOF-MS. Briefly, an aliquot of VLPs (10 μL from a 2 mg/mL stock) was denatured by treatment with DTT and urea (0.2 M final concentration each), followed by a brief incubation at room temperature (< 5 m). Denatured particles were acidified by addition of TFA (0.1% final concentration), and subjected to solid-phase extraction by use of C4 ZipTips (EMD Millipore) according to the manufacturer's instructions.

The desalted samples were eluted using 10 μL of 70:30/acetonitrile:water in 0.1% TFA and immediately loaded on a 6230B time-of-flight LC/MS (Agilent). 5-10 μL of each sample was injected onto a Poroshell 300SB-C3 LC column (Agilent), run in positive mode, using the following solvents and run conditions:
Solvent A: 0.1% formic acid in HPLC-grade water; Solvent B: 100% acetonitrile. Initial condition: 10% B, 0-2 minutes; 10-50% B, 2-5 minutes; 50-95% B, 5-10 minutes; 1 minute re-equilibration at 10% B.

Data were acquired from 250-2500 m/z using the following TOF source parameters: drying gas 325 °C and flow 13 L/min; nebulizer 40 psi; sheath gas 325 °C and flow 12 L/min; capillary 4000 V; nozzle 1000 V; fragmentor 250 V; skimmer 60 V. Raw MS data were analyzed using

MassHunter BioConfirm software (Agilent) and deconvoluted using a maximum entropy algorithm, which yielded the masses of the coat protein subunits (the modified PP7-PP7 proteins). Under the assumption that particle modification does not significantly impact ionization efficiency, we calculated the apparent extent of protein modification following each chemical conjugation step.

**Bioconjugation**

**Preparation of fluorescently labeled azide-modified PP7-PP7 VLPs**. To a solution of PP7-PP7 VLPs (1 mL from 6 mg/mL stock in 0.1 M potassium phosphate buffer; 0.21 μmol PP7-PP7 subunit), was added 14 μL 25 mM NHS-AF647 (0.35 μmol NHS, 1.67 equivalents to PP7-PP7 subunit). The reaction mixture was wrapped in foil and placed on rotating incubator at room temperature. After 2 hours, a solution of N-hydroxysuccinimidyl ester-containing azide (150 μL from 250 mM stock in DMSO, freshly prepared; 37.5 μmol, ~178.6 equivalents to PP7-PP7 subunit) was added, and the reaction mixture was further incubated at room temperature. After two additional hours, particles were purified by PD-10 desalting column (Cytiva), followed by centrifugal filtration using an Amicon Ultra-4 100k MW cut-off device (EMD Millipore). Protein recovery was determined via a Bradford assay against BSA standards and was typically ~70-75%. The extent of particle acylation was determined by LC-ESI-TOF-MS analysis, as described below (~900-1100 azides per VLP; 7.5-9.2 azides per subunit). The extent of fluorophore labeling was determined by UV-Vis, comparing the AF647 concentration against the particle concentration (typically ~30-40 AF647/VLP).

**Preparation of peptide and ligand-labeled PP7-PP7 VLPs**. OvaI or OvaII peptides were clicked to separate particles. First, we determined the relative efficiency of peptide attachment for OvaI versus OvaII peptides. PP7-PP7-(azide)$_{930}$ VLPs (8.5 μL from 4.7 mg/mL stock; ~1.2 mM in VLP-displayed azide) were combined with a premixed solution of 5:1 tris((1-hydroxy-propyl-1H-1,2,3-triazol-4-yl)methyl)amine (THPTA):Cu (0.57 μL from stock of 400 mM THPTA:80 mM Cu; 45.6 nmol, ~4.5 equivalents to VLP-azide), an aliquot of aminoguanidine (0.8 μL from 0.5 M stock in water), an aliquot of sodium ascorbate (0.8 μL from 0.5 M stock in water), and an aliquot of either OvaI or OvaII alkyne (from a 5 mM stock in DMSO). The amount of peptide-alkyne in each reaction was varied from 0.8 μL (4 nmol, ~0.4 equivalents to VLP-azide), to 2 μL (10 nmol, ~1.0 equivalents to VLP-azide), to establish concentrations needed to load approximately 90-120 peptides per capsid. Approximately 0.4-0.5 equivalents of OvaI and 0.5-0.6 equivalents of OvaII (relative to VLP-azide) were used henceforth.
To scale up reactions for *in vitro* and *in vivo* experiments, a solution of PP7-PP7-azide (280 μL from 5 mg/mL stock in 0.1 M potassium phosphate buffer, ~350 nmol azide, 1 equivalent azide) was combined with a premixed solution of 5:1/THPTA:Cu (19.8 μL from stock of 400 mM THPTA:80 mM Cu; ~1.58 μmol, ~4.5 equivalents to VLP-azide), an aliquot of aminoguanidine (19.8 μL from 0.5 M stock in water), an aliquot of sodium ascorbate (19.8 μL from 0.5 M stock in water), and an aliquot of OvaI or OvaII alkyne (32 μL from a 5 mM stock of OvaI, 0.45 equivalents relative to VLP-azide; or 42 μL from a 5 mM stock of OvaII, 0.6 equivalents relative to VLP-azide). Reactions were capped and mixed by gentle inversion, then incubated at 50 °C with rotation. After 10 minutes, a solution of mannose alkyne (34 μL from 25 mM stock in DMSO; 0.85 μmol, ~2.4 equivalents to VLP-azide) was added to each reaction. Reactions were incubated at 50 °C with rotation for an additional 50 minutes. Modified particles were recovered using a PD-10 desalting column, followed by centrifugal filtration using Amicon Ultra 30 kDa

MWCO filters. Protein recovery was determined via a Bradford assay (typically ~70% recovery) and particles were characterized as described above to determine particle stability, purity, and the extent of modification. Levels of endotoxin contamination were found to be less than 0.1 EU/mL using Pierce LAL Chromogenic Endotoxin Quantitation Kit (Thermo Fisher).

**Cell lines and mouse strains**

Adult human peripheral blood was acquired from Research Blood Components, LLC (Boston, MA, USA) and Massachusetts General Hospital Blood Donor Center (Boston, MA, USA) under an institutional review board (IRB)-approved protocol. Monocytes were isolated from whole blood by negative selection using RosetteSep human monocyte enrichment cocktail following the manufacturer's recommendations (StemCell Technologies). Using density gradient medium (Lymphoprep), monocytes were isolated and differentiated into immature DCs in the presence of GM-CSF (100 ng/mL) and IL-4 (50 ng/mL) (R&D Systems) in CellGenix GMP DC media (Sartorius) for 6 to 7 days. B16F10-OVA cells were kindly provided by K. Dane Wittrup at MIT. B16F10-OVA cells were cultured in complete medium (DMEM, 10% FBS and 100 U $ml^{-1}$ penicillin G sodium and 100 μg $ml^{-1}$ streptomycin) at 37 °C in 5% $CO_2$.

Female C57BL/6 mice were obtained from Jackson Laboratory and used at 8 weeks of age. Experiments were performed in specific pathogen-free animal facilities at the MIT Koch Institute for Integrative Cancer Research. All mouse studies were performed according to institutional and National Institutes of Health guidelines for humane animal use and in accordance with the Association for Assessment and Accreditation of Laboratory Animal Care. All protocols were approved by the Institutional Animal Care and Use Committee at MIT.

**VLP Internalization**

The moDCs were suspended in phosphate-buffered saline (PBS) (pH 7.4) supplemented with 1 mM $CaCl_2$ and 0.5 mM $MgCl_2$ at $1.0 \times 10^6$ cells/mL. Cells were stimulated with 8 nM AF647-functionalized VLPs at 37 °C for the indicated time points. To inhibit the uptake of VLPs, moDCs were pretreated with 10 μg/mL mouse anti-human CD209 antibody clone DCN46 (BD Biosciences), mouse anti-human CD206 antibody clone 19.2 (BD Biosciences), human Dectin-2/CLEC6A antibody clone # 545925 (R&D Systems), or human CLEC4E antibody clone # 2455C (R&D Systems) for 20 min on ice followed by incubation with the VLPs at 37 °C for 15 min. Cells were washed in PBS + 0.1% BSA and treated with human Fc block (BD Biosciences) for 15 min at rt followed by staining with Ghost Red 780 (Cytek Biosciences) and BV785 anti-human CD11c antibody clone 3.9 (Biolegend). The extent of VLP internalization was measured by flow cytometry (BD FACSymphony A3). Data were analyzed using FlowJo software (v.10.9) utilizing the geometric mean fluorescence intensity (MFI) to quantify VLP internalization on $CD11c^+$ moDCs.

**Microscopy**

To assess VLP trafficking, moDCs ($0.5 \times 10^6$ cells/mL) were incubated with Cy3 human transferrin (Jackson ImmunoResearch Laboratories Inc.) for 20 min at 37 °C to label the endosomes, followed by incubation with 8 nM VLPs for 30 min at 37 °C. To assess DC-SIGN and TLR7 localization, moDCs ($0.5 \times 10^6$ cells/mL) were incubated with 8 nM VLPs for 30 min at 37 °C. Cells were washed and permeabilized using the fixation/permeabilization kit following the manufacturer's recommendations (BD Biosciences). Permeabilized cells were then washed and stained for 30 min at 4 °C with AF405 human DC-SIGN antibody clone # 120507 (R&D Systems) and AF488 human TLR7 antibody clone # 533707 (R&D Systems). Following staining, cells were washed and transferred to 96 well glass bottom plate with #1.5 cover glass (Cellvis). Cells were imaged using a Zeiss Cell Discoverer 7 + LSM900 confocal microscope (Carl Zeiss Microscopy USA), with a 50× 1.2NA water objective and 60 pinhole and analyzed on the Zen Microscopy Software (Carl Zeiss Microscopy USA).

**DC Activation**
The moDCs were suspended in CellGenix GMP DC media at $1.0 \times 10^6$ cells/mL. Cells were stimulated with 8 nM VLPs at 37 °C. After 24 h, cells were washed in PBS + 0.1% BSA and treated with human Fc block (BD Biosciences) for 15 min at rt followed by staining with Ghost Red 780 (Cytek Biosciences) and the anti-human Ab cocktail: BV785 anti-human CD11c antibody clone 3.9 (Biolegend), PE anti-human CD80 antibody clone 2D10 (Biolegend), BV421 anti-human CD83 antibody clone HB15e (Biolegend), BV605 anti-human CD86 antibody clone BU63 (Biolegend), and BUV395 mouse anti-human CD40 antibody clone 5C3 (BD Biosciences). Cells were washed in PBS + 0.1% BSA and analyzed by flow cytometry (BD FACSymphony A3). Data were analyzed using FlowJo software (v.10.9) utilizing the geometric mean fluorescence intensity (MFI) to quantify receptor expression on $CD11c^+$ moDCs.

**RNA Sequencing**
For RNA-Seq analyses, moDCs were treated with 8 nM VLPs for 6 h at 37 °C. Total RNA was isolated using NucleoSpin RNA kits according to the manufacturer's recommendations (Takara). RNA was purified using RNA SPRIbeads before being evaluated using an Agilent Fragment Analyzer. mRNA from 40ng of RNA was isolated and used to prepare Illumina libraries as previously described.[5] Libraries were sequenced on an Illumina NextSeq500 with 40nt paired end reads. FASTQ files were mapped to the reference human genome GRCh38 using STAR.[6] Samples with low alignment rates and/or unique alignment issues were excluded from further downstream analyses. Differential expression analyses were performed in R using the DESeq2 package,[7] with a multifactor design accounting for biological replicate (donor) and condition. For gene signature module scoring raw counts from alignment were log-normalized and scaled in Seurat and then the AddModuleScore function in Seurat was implemented. To calculate a gene module score for TLR7 activation, a gene signature was obtained from the literature.[8] For visualization on a heatmap, z-scores were calculated on data normalized using variance stabilizing transformation implemented in DESeq2. GSEA was performed using fGSEA algorithm to compute normalized enrichment scores and associated *P* values using the Hallmark gene signatures from mSigDB.[9]

**Cytokine Analysis**
The moDCs were stimulated with 8 nM VLPs for 32 h at 37 °C. The supernatant from these samples was then collected and analyzed using the Legendplex Human Anti-Virus Response Panel (Biolegend) according to the manufacturer's recommendations and analyzed by flow cytometry (Attune NxT). Cytokine concentrations were calculated against standard curves using Qognit software (Biolegend).

**Biodistribution**

Animals were injected s.c. with 50 μg AF647-functionalized VLPs in PBS. The animals were euthanized at the indicated timepoints. To assess AF647-VLP biodistribution, animals were sacrificed at the indicated timepoints and tissues, including LNs, spleen, kidneys, liver, heart, and lungs, were excised and imaged at 720 nm with 640 nm excitation, with an exposure time of 0.1 s. Fluorescent imaging was performed with an IVIS Spectrum instrument (Perkin Elmer). Radiant efficiency ($p\ s^{-1}\ sr^{-1}\ \mu W^{-1}$) was evaluated for each experiment and treatment group using elliptical regions of interest in Living Image Software (v.4.7.2) (Perkin Elmer). To assess cellular uptake in the LNs, mice were euthanized after 24 h and LN cells were excised and stained as previously reported.[10] Briefly, LNs were incubated with 100 μl of 10 mg/mL collagenase D (Sigma Aldrich) at 37 °C for 1 h. The LNs were then mechanically disrupted on a 40 μm cell strainer and cells were plated at a concentration of $1 \times 10^6$ cells/mL. Cells were blocked with Fc block rat anti-mouse CD16/CD32 (BD Biosciences) followed by Live Dead staining with Ghost Red 780 (Cytek Biosciences) and stained using BV421 hamster anti-mouse CD11c antibody clone HL3 (BD Biosciences). Cells were washed with PBS + 0.1% BSA and analyzed by flow cytometry (BD FACSymphony A3). DCs were gated as $CD11c^+$ and the extent of AF647-VLP uptake was quantified as a geometric mean fluorescence intensity (MFI) using FlowJo software (v.10.9).

**Mouse Immunization and Sampling**

C57BL/6 mice were injected subcutaneously with $5 \times 10^5$ B16F10-OVA melanoma cells on the right-side flank on day 0. Mice were then randomly divided into different cohorts and immunized s.c. with 50 μg VLPs and 10 μg 2'3'-cGAMP VacciGrade (Invivogen) as an adjuvant in PBS and i.p. with 100 μg InVivoPlus anti-mouse PD-1 (BioXCell) on days 3, 9, and 15. Tumor measurements were taken using calipers every 3 days after tumor inoculation. Serum samples were collected on day 21 via cardiac puncture, processed using a serum separator blood collection tube (BD Biosciences), and stored at -20 °C. For intracellular cytokine staining, splenocytes were isolated at day 21 and processed for flow cytometry as described below.

**Intracellular Cytokine Staining**

Splenocytes were isolated from mice following a protocol described previously.[3] Briefly, spleens were excised and mechanically disrupted on a 70 μm cell strainer. Cells were incubated with RBC lysis buffer (Invitrogen) for 2 min and washed with RPMI medium. Splenocytes were resuspended at a concentration of $2 \times 10^6$ cells/well in RPMI medium supplemented with 10% FBS and cultured in U-bottom tissue culture plates (Nunc, Denmark) in the presence of OVA(323−339) peptide (5 μg/mL) (Invivogen), OVA(257−264) peptide (5 μg/mL) (Invivogen), or phorbol myristate acetate−ionomycin cocktail (Invitrogen) as a positive control. Unstimulated splenocytes were included as a negative control. Splenocytes and antigens were incubated for 1 h at 37 °C in 5% $CO_2$. After 1 h, brefeldin A (BD Biosciences) was added and cell cultures were incubated for an additional 5 h. Following the 6 h stimulation, cells were washed with PBS and stained with Ghost Dye Red 780 (Cytek Biosciences). Cells were then blocked with Fc block rat anti-mouse CD16/CD32 clone 2.4G2 (BD Biosciences) and stained with cell surface monoclonal antibodies: BV421 anti-mouse CD3 antibody clone 17A2 (Biolegend), APC anti-mouse CD4 antibody clone RM4-5 (Biolegend), and BUV395 rat anti-mouse CD8a antibody clone 53-6.7 (BD Biosciences). Following surface staining, cells were permeabilized using the fixation/permeabilization kit following the manufacturer's recommendations (BD Biosciences).

Permeabilized cells were then washed and stained for 30 min at 4 °C with Alexa Fluor 488 anti-mouse TNF-α antibody clone MP6-XT22 (Biolegend) and PE anti-mouse IFN-γ antibody clone XMG1.2 (Biolegend). Following staining, cells were washed with PBS + 0.1% BSA and analyzed by flow cytometry (BD FACSymphony A3).

**Detection of Antibody Response in Serum**

Anti-OVA(323-339) IgG antibody response was measured in serum by enzyme-linked immunosorbent assay (ELISA). MaxiSorp plates (Thermo Fisher) were coated with streptavidin (10 μg/mL) (Sigma Aldrich) in PBS at 4 °C overnight and washed three times with PBST (PBS + 0.05% Tween 20). Plates were blocked with 1× casein blocking buffer (Sigma Aldrich) in PBS at rt for 1 h and washed three times with PBST. Plates were then incubated with biotin-labeled OVA(323-339) peptide (5 μg/mL in blocking buffer) (AnaSpec) at rt for 1 h and washed three times with PBST. Serum (2× dilution series from 1:100 to 1:204800 in blocking buffer) were added to the plate and incubated at rt for 2 h followed by three washes with PBST. The presence of OVA(323-339)-specific antibodies was detected using horseradish peroxidase-conjugated goat anti-mouse IgG, IgG1, IgG2b, IgG2c, IgG3 (Southern Biotech Assoc Inc), and IgM (Invitrogen) antibodies (diluted 1:2000 in blocking buffer). Rabbit anti-chicken OVA(323-339) IgG (Alpha Diagnostic International) was used as a control and labeled with goat anti-rabbit IgG-HRP (Southern Biotech Assoc Inc). After 2 h of incubation at rt, the plates were washed four times with PBST and developed with 3,3′,5,5′-tetramethylbenzidine (TMB) Liquid Substrate System (Sigma Aldrich) for 15 minutes and quenched with 2M $H_2SO_4$. The absorbance at 450 nm was determined using a SpectraMax M5 microplate reader (Molecular Devices, Sunnyvale, CA). The endpoint titers were calculated using GraphPad Prism 10 (GraphPad Software, Inc., La Jolla, CA, USA) to be the maximum serum dilution that displayed an optical density (OD) value of ≥0.3 using a nonlinear regression of the sigmoidal curve (antisera dilution against OD at 450 nm) followed by a four-parameter logistic curve analysis.

**Statistical Analysis**

Statistical analyses were performed using GraphPad Prism 10 (GraphPad Software, Inc., La Jolla, CA, USA). Wilcoxon matched-paired signed rank test and one-way or two-way ANOVA with multiple comparisons tests were performed as indicated. All reported P values were two tailed, and statistical significance was defined as a P value of $<0.05$.

**Data Availability**

The RNA-seq data are available in the NCBI Gene Expression Omnibus (GEO) under accession no. GSE251818. The authors declare that other data supporting the findings of this study are available within this article and its supplementary information; all additional data are available from the corresponding author upon reasonable request.

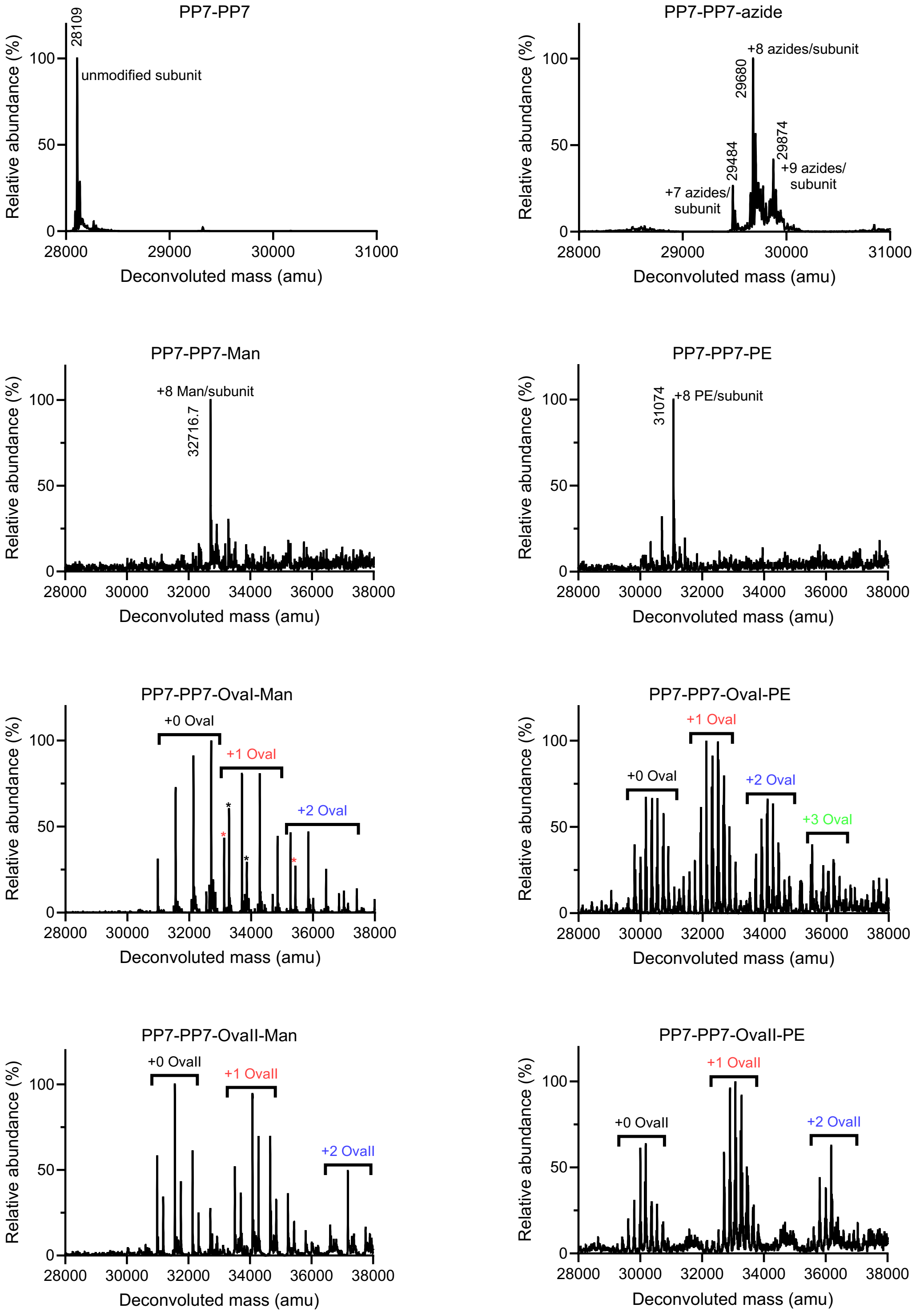


**Fig. S1 | Deconvoluted ESI-TOF HRMS for PP7-PP7 conjugates.** The average number of azides, glycomimetic ligands, or peptides installed onto each PP7-PP7 subunit are indicated; each particle is composed of 120 copies of subunits.

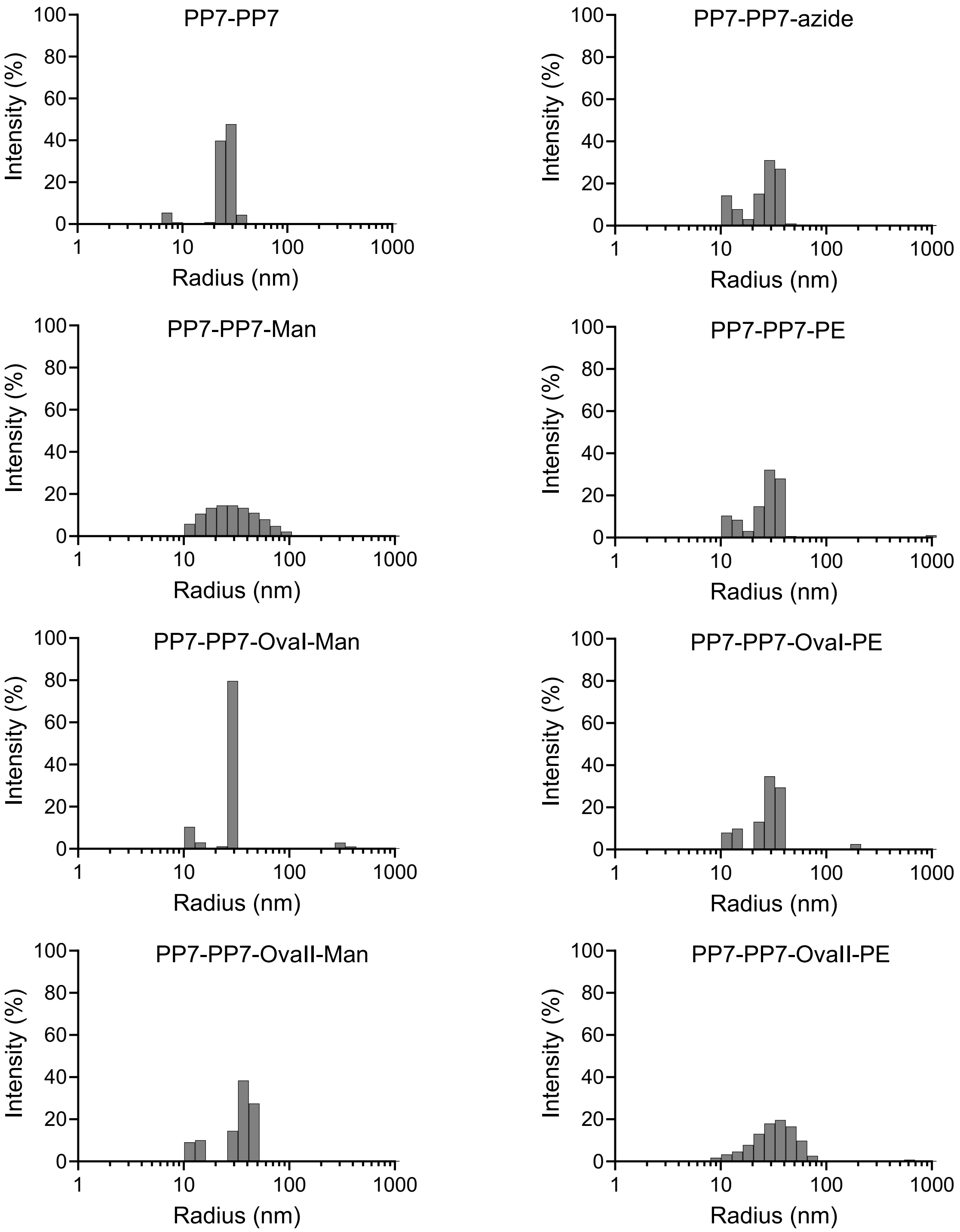


**Fig. S2 | Dynamic light scattering analysis of PP7-PP7 conjugates.** Protein samples were diluted in 0.1 M potassium phosphate buffer (50 µL, final conc: 0.1 mg/mL) and the hydrodynamic radius of each sample was measured using a Dynapro plate reader (approx. 3,000,000 counts per second, 30 measurements per sample).

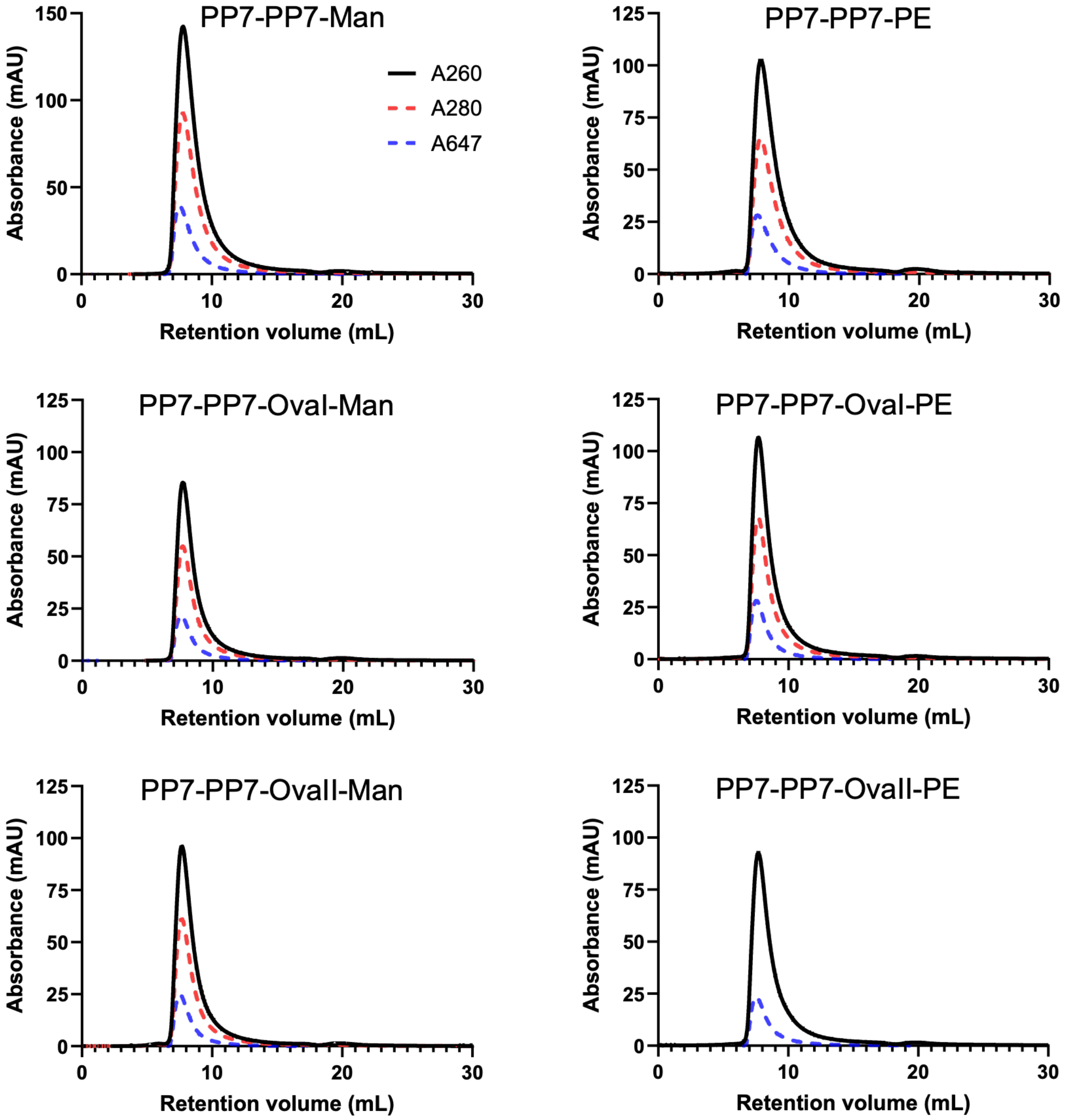


**Fig. S3 | FPLC and UV-Vis characterization of PP7-PP7 VLP conjugates.** FPLC profile of PP7-PP7 conjugates (100 µL injection of 0.1 mg/mL VLPs); black solid line = A260, red dashed line = A280, blue dashed line = A647.

**Table S1 |** PP7-PP7 conjugate vaccines, all bearing approximately ~30-40 AlexaFluor647 dyes per particle. a) Average number of attached ligands per particle. b) Average number of attached peptides per particle.

| Entry | Particle | Ligand | #Lig[a] | Peptide | #Pep[b] | r [nm] |
|---|---|---|---|---|---|---|
| **1** | PP7-PP7-Man | Man | 960-1080 | - | - | 26.2 |
| **2** | PP7-PP7-Man-OvaI | Man | 850-950 | OvaI | 80-120 | 27.4 |
| **3** | PP7-PP7-Man-OvaII | Man | 850-950 | OvaII | 80-120 | 29.3 |
| **4** | PP7-PP7-PE | PE | 960-1080 | - | - | 24.9 |
| **5** | PP7-PP7-PE-OvaI | PE | 850-950 | OvaI | 80-120 | 26.6 |
| **6** | PP7-PP7-PE-OvaII | PE | 850-950 | OvaII | 80-120 | 30.0 |
| **7** | PP7-PP7-azide | Azide | 960-1080 | - | - | 23.4 |

**Table S2 |** Sequence of particle-displayed peptides used. a) Peptides consisted of a propargyl-linker (italicized), a Cathepsin D-cleavable motif (underlined), and the OVA(257-264) or OVA(323-339) epitopes (bolded). b) Mass of Ova peptide (bolded sequence) delivered per 100 µg of VLPs, each bearing ~80-120 copies of peptide.

| Entry | Peptide | Sequence[a] | Typical Ova loading (µg Ova per 100 ug particle)[b] |
|---|---|---|---|
| **1** | OvaI | $^{11}$*GSGS*DGSPLEF**SIINFEKL** | 2.6-4.0 |
| **2** | OvaII | $^{11}$*GSGS*DGSPLEF**ISQAVHAAHAEINEAGR** | 4.6-6.2 |

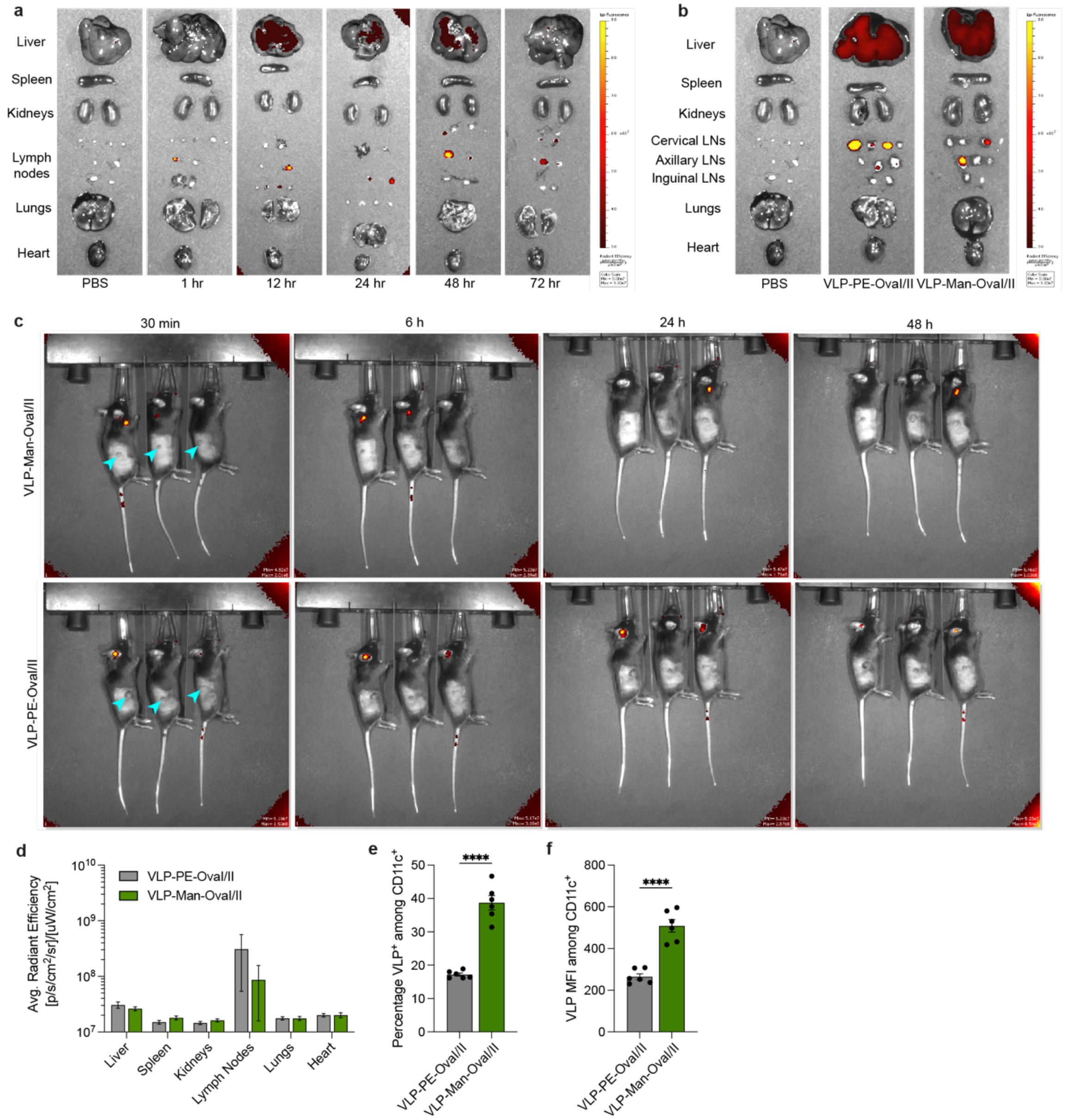


**Fig. S4 | VLPs traffic to lymph nodes for uptake by LN-resident immune cells. a,** C57BL/6 mice were immunized with phosphate-buffered saline (PBS) or VLP-PE-OvaI/II and organs were isolated for quantification by IVIS imaging at indicated timepoints **b,** C57BL/6 mice were immunized with PBS or AF647-labeled VLPs and organs were isolated for quantification by IVIS imaging at 24 h. **c,** C57BL/6 mice were inoculated with tumors for 7 days followed by immunization with AF647-labeled VLPs. VLP trafficking to the tumor was assessed by IVIS imaging over the course of 48 h. Arrowheads indicate flank tumor site. **d,** Quantification of average radiant efficiency within each organ of interest at 24 h (value for PBS control tissue ≈ 1.4-2×$10^7$). **e,f,** LN-resident $CD11c^+$ DCs were analyzed for AF647-labeled VLP uptake by flow cytometry. Data represent mean  s.e.m. from a representative experiment (n=3 (**d**) and n=6 (**e-f**)). Statistical analysis was performed by unpaired two-tailed Student's *t*-test. ****P<0.0001.
